# CROP: A Retromer-PROPPIN complex mediating membrane fission in the endo-lysosomal system

**DOI:** 10.1101/2021.09.06.459059

**Authors:** Thibault Courtellemont, Maria Giovanna De Leo, Navin Gopaldass, Andreas Mayer

## Abstract

Endo-lysosomal compartments exchange proteins by fusing, fissioning, and through endosomal transport carriers. Thereby, they sort many plasma membrane receptors and transporters and control cellular signaling and metabolism. How the membrane fission events are catalyzed is poorly understood. Here, we identify the novel CROP complex as a factor acting at this step. CROP joins members of two protein families: the peripheral subunits of retromer, a coat forming endosomal transport carriers, and membrane inserting PROPPINs. Integration into CROP potentiates the membrane fission activity of the PROPPIN Atg18 on synthetic liposomes and confers strong preference for binding PI(3,5)P_2_, a phosphoinositide required for membrane fission activity. Disrupting CROP blocks fragmentation of lysosome-like yeast vacuoles *in vivo*. CROP-deficient mammalian endosomes accumulate micrometer-long tubules and fail to export cargo, suggesting that carriers attempt to form but cannot separate from these organelles. PROPPINs compete for retromer binding with the SNX proteins, which recruit retromer to the membrane during the formation of endosomal carriers. Transition from retromer-SNX complexes to retromer-PROPPIN complexes might hence switch retromer activities from cargo capture to membrane fission.

## Introduction

Endo-lysosomal compartments play a major role in controlling the abundance of most plasma membrane transporters and receptors and therefore have a key role in defining the communication, interaction, and transport capacities of a cell. They receive proteins from the cell surface or from the Golgi and sort them either for recycling back to these compartments, or for transfer to lysosomes, where many of them become degraded (Cullen & Steinberg, 2018; Ma & Burd, 2019; Seaman, 2019). Endo-lysosomal compartments exchange proteins and lipids through homo- and heterotypic fusion and fission events and through tubular-vesicular carriers. These structures can be formed by a variety of membrane coats, which can recruit cargo into them, such as retromer, retriever, CCC or ESCPE (Cullen & Steinberg, 2018).

Retromer is a conserved coat that consists of a peripheral part and of a lipid-interacting part. In yeast, retromer has been discovered as a stable entity that could be dissociated into two subcomplexes, consisting of the sorting nexins Vps5 and Vps17 (SNX complex) and of a complex of Vps26, Vps29 and Vps35 (CSC complex) (Seaman *et al*, 1998). SNX binds membranes via PX domains, which recognize phosphatidylinositol-3-phosphate (PI3P) (Burda *et al*, 2002), and via BAR domains, which preferentially associate with highly curved bilayers. The Vps26/29/35 complex by itself shows only weak affinity for the membrane and requires SNX for recruitment. Retromer associates with cargo and numerous other factors, which are important for the formation of the transport carriers and/or their fission from the membrane. These include components of the Rab-GTPase system (Seaman *et al*, 2009; Rojas *et al*, 2008; Da Jia *et al*, 2016; Balderhaar *et al*, 2010; Liu *et al*, 2012; Purushothaman & Ungermann, 2018), the actin-regulating WASH complex (Jia *et al*, 2012; Harbour *et al*, 2012; Chen *et al*, 2019; Derivery *et al*, 2009; Gomez & Billadeau, 2009; Phillips-Krawczak *et al*, 2015; Rojas *et al*, 2007; Temkin *et al*, 2011; Lucas *et al*, 2016), or EHD1, and ATPase with structural similarities to dynamins (Daumke *et al*, 2007; Gokool *et al*, 2007).

Structural analyses of retromer shed light onto its mode of action (Hierro *et al*, 2007; Lucas *et al*, 2016; Kovtun *et al*, 2018; Kendall *et al*, 2020; Collins *et al*, 2008; 2005; Purushothaman *et al*, 2017). These studies begin to elucidate cargo recognition, the organization of the subunits on the membrane and the way in which their association promotes membrane tubulation. The subsequent step of detaching the carrier from the donor membrane is promoted by numerous protein factors, but their mechanism of action in the context of retromer is still poorly understood: the actin-regulating WASH complex could enhance fission by increasing membrane tension and friction (Bar-Ziv *et al*, 1999; Markin *et al*, 1999; Derivery *et al*, 2009; Gomez & Billadeau, 2009; Phillips-Krawczak *et al*, 2015; Simunovic *et al*, 2017). Mechanochemical factors, such as the dynamin-like GTPase Vps1 or ATPases of the EHD family, might constrict the membranes (Chi *et al*, 2014; Deo *et al*, 2018). Fission appears to be favored also by contact of the endosomal transport carriers to ER membranes (Rowland *et al*, 2014). Recently, we described fission activity of the PROPPIN Atg18 on synthetic liposomes (Gopaldass *et al*, 2017) and showed that its human homolog WIPI1 is required for protein exit from endosomes (De Leo *et al*, 2021).

PROPPINs form a protein family that is present with multiple isoforms in eukaryotic cells from yeast to men (Dove *et al*, 2004). Baker’s yeast expresses three isoforms, Atg18, Atg21 and Hsv2, and mammalian cells express four genes (WIPI1 through 4). PROPPINs bind phosphoinositides phosphorylated on the 3- and/or 5-position and support the assembly of the autophagic machinery on phagophores (Vicinanza *et al*, 2015; Baskaran *et al*, 2012; Krick *et al*, 2012; Liang *et al*, 2019; Proikas-Cezanne *et al*, 2004; Watanabe *et al*, 2012). In autophagy, WIPI proteins interact with and recruit key factors of this machinery, such as Atg16L1, Atg5, Atg12 and Atg2. They also participate in autophagic signaling and promote the interaction of the isolation membrane with the ER (Stromhaug *et al*, 2004; Obara *et al*, 2008; Polson *et al*, 2010; Dooley *et al*, 2014; Proikas-Cezanne *et al*, 2015; Bakula *et al*, 2017; Itakura & Mizushima, 2010; Lu *et al*, 2011) (Chowdhury *et al*, 2018; Stanga *et al*, 2019; Otomo *et al*, 2018; Lei *et al*, 2020). The autophagic functions of PROPPINs depend on phosphatidylinositol-3-phosphate (PI3P) or phosphatidylinositol-5-phosphate (PI5P) (Vicinanza *et al*, 2015; Baskaran *et al*, 2012; Krick *et al*, 2012; Liang *et al*, 2019; Proikas-Cezanne *et al*, 2004; Watanabe *et al*, 2012).

PROPPINs are, however, not restricted to the sites of autophagosome formation. They show strong enrichment on endo-lysosomal organelles, where they reduce the size of endosomes and influence the distribution of protein between the endosomes and the Golgi or vacuoles (Jeffries *et al*, 2004; Dove *et al*, 2004). The yeast PROPPIN Atg18 promotes the division of the vacuole into smaller fragments (Dove *et al*, 2004; Efe *et al*, 2007; Gopaldass *et al*, 2017; Zieger & Mayer, 2012; Michaillat *et al*, 2012). The human PROPPIN WIPI1 is required in multiple protein exit pathways from endosomes, which transfer proteins to the plasma membrane, to the Golgi, or to lysosomes. Here, it promotes the PI3P-dependent formation of endosomal transport carriers and their phosphatidylinositol-3,5-bisphosphate (PI(3,5)P_2_) dependent fission from endosomes (De Leo *et al*, 2021). The endosomal and autophagic functions of Atg18 and WIPI1 can be differentiated through several molecular features and interactors, which are relevant only for one of the two processes (Gopaldass *et al*, 2017; De Leo *et al*, 2021).

When incubated with synthetic giant unilamellar vesicles (GUVs) at micromolar concentrations, pure recombinant Atg18 suffices to tubulate these membranes and divide them into small vesicles, underlining its potential as a membrane fission protein (Gopaldass *et al*, 2017). These in vitro assays revealed that fission depends on a hydrophobic loop of Atg18, which can fold into an amphipathic helix when brought in contact with the membrane. The amphipathic helix is conserved in other PROPPINs and it is essential for membrane fission on mammalian endosomes as well as on yeast vacuoles in vivo (Gopaldass *et al*, 2017; De Leo *et al*, 2021). It was proposed that loop insertion promotes fission by increasing membrane curvature. This effect may be amplified by oligomerisation of Atg18, which could not be induced through PI3P, but only through PI(3,5)P_2_ (Gopaldass *et al*, 2017; Scacioc *et al*, 2017). PROPPINs are thus likely to be the effector proteins of PI(3,5)P_2_, the lipid that is necessary to drive fission on a variety of endo-lysosomal compartments (McCartney *et al*, 2014). Although there appears to be specificity for PI(3,5)P_2_ in these functional terms, purified PROPPINs bind PI3P, PI5P and PI(3,5)P_2_ fairly promiscuously (Vicinanza *et al*, 2015; Baskaran *et al*, 2012; Krick *et al*, 2012; Liang *et al*, 2019; Proikas-Cezanne *et al*, 2004; Watanabe *et al*, 2012; Busse *et al*, 2015).

While these in vitro experiments clearly demonstrated the potential of PROPPINs to promote membrane fission, they could not resolve whether perform this function alone in the cellular context. Using the yeast PROPPIN Atg18 we hence began to search for interactors that might participate in the Atg18-dependent fission of yeast vacuoles. This led us to discover a novel complex, which we term CROP. CROP integrates Atg18 with a part of the endosome- and vacuole-associated retromer complex to generate a membrane fission device of much higher potency. We studied its activity in yeast, in human cells and on liposomes.

## Results

### Atg18 forms a complex with Vps26/29/35

Seeking proteins that might cooperate with the PROPPIN Atg18 in driving membrane fission on yeast vacuoles, we affinity-purified a FLAG-tagged Atg18 (Atg18^Gly6-FLAG3^) expressed under its native promoter in yeast. Associated proteins were quantified by mass spectrometry, using SILAC (stable isotope labelling by amino acids in cell culture) (Fig. 1a). 11 proteins were significantly interacting with Atg18 (Suppl. Table 1a). The most abundant interactor in normal rich medium was Atg2, a protein binding Atg18 during autophagosome formation (Obara *et al*, 2008). Other Atg18 interactors, such as the phosphatase Sit4, which is required for vacuole fission (Michaillat *et al*, 2012; Michaillat & Mayer, 2013), and its regulatory subunits Cdc55 and Sap155, were detected in extracts from cells in which vacuole fission had been triggered by a moderate osmotic shock (Zieger & Mayer, 2012) (Fig. 1b). Also three proteins from the retromer complex (Vps26, Vps35, and Vps29) bound Atg18 more strongly upon triggering vacuole fission. They form a stable subcomplex, called cargo-selective complex (CSC) (Seaman *et al*, 1998). Our mass spectrometry analysis had not identified any peptides from Vps5 or Vps17, the phosphatidylinositol-3-phosphate (PI3P)-interacting sorting nexins (SNX complex) that recruit CSC to the membrane (Seaman *et al*, 1998).

**Figure 1:**
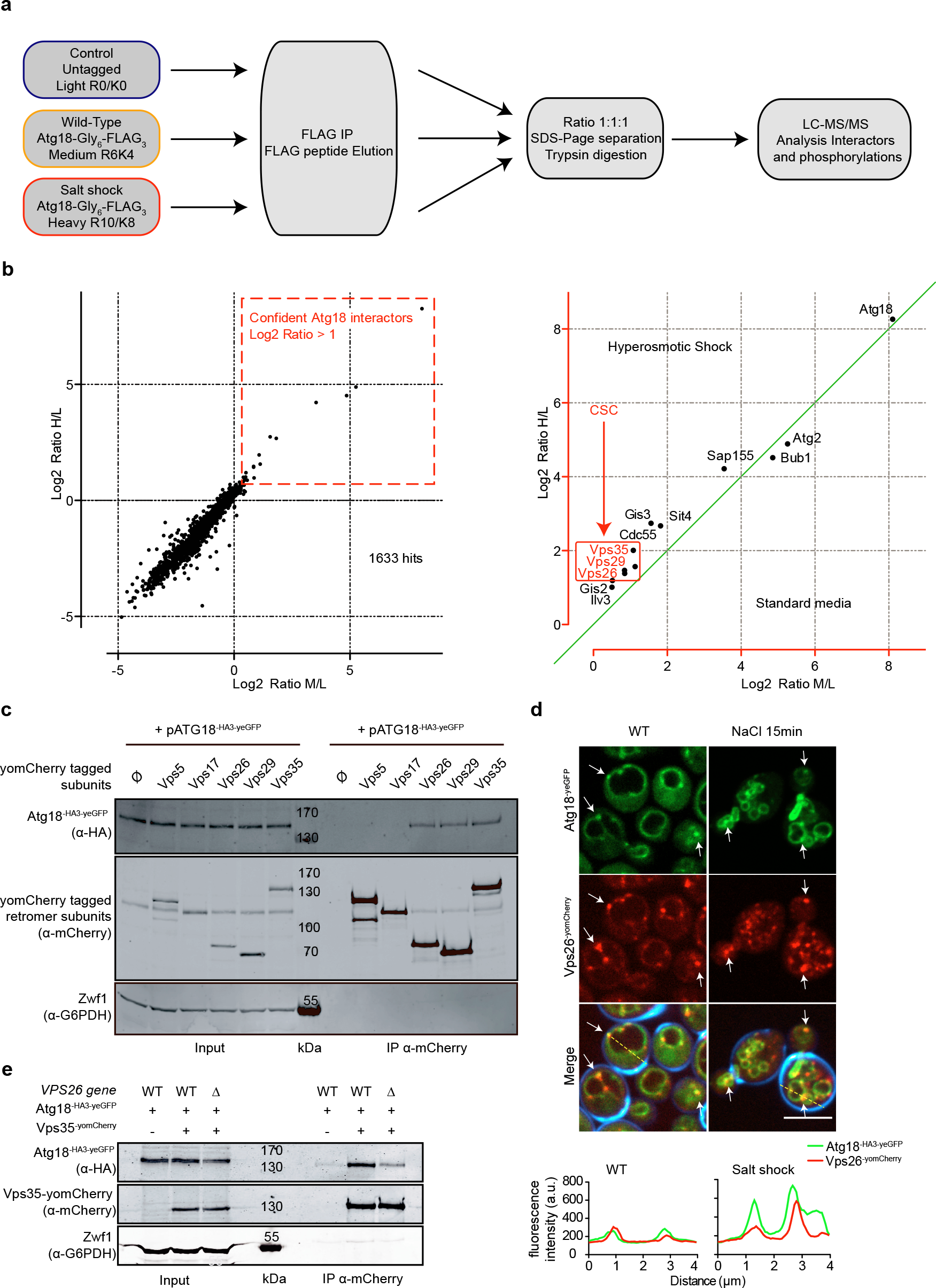
Atg18 interacts with Vps26, Vps29 and Vps35. **a**, Procedure of the Triplex SILAC approach. **b**, Scatter plot of the log2 distribution of Atg18 partners identified by SILAC mass spectrometry. The abscissa shows log2 ratios of peptides found in standard media relative to the non-tagged negative control (medium/light; M/L). The ordinate shows log2 ratios of peptides found in salt shocked cells, relative to the non-tagged negative control (heavy/light; H/L). **c**, Interaction of Atg18 with retromer subunits. Genomically tagged yomCherry-fusions of each retromer subunit were expressed in *SEY6210 atg18*Δ, *atg21*Δ, *hsv2*Δ cells, together with a plasmid expressing Atg18^HA3yeGFP^. Cell lysates were subjected to immunoprecipitation using RFP-Trap magnetic beads and analyzed by SDS-PAGE and Western blotting against the indicated proteins. **d**, Salt-induced vacuole fragmentation. Live cell confocal imaging of Atg18^HA3-yeGFP^ and Vps26^yomCherry^ before and after a mild salt shock with 0.5 M NaCl for 15 minutes. Calcofluor white used to stain the cell walls is only shown in the merge. Scale bar: 5 µm. A line scan along the dashed yellow line shown in d. **e**, Requirement of Vps26 for the Atg18-Vps35 interaction. Genomically tagged Vps35^yomCherry^ was pulled down from lysates of *SEY6210 WT* or *SEY6210 vps26*Δ strains. Adsorbed proteins were analyzed by SDS-PAGE and Western blotting using the antibodies indicated in brackets.

We tested the CSC-Atg18 interaction through immunoadsorption. To this end, we tagged all retromer subunits individually with yomCherry and expressed them at their genomic locus, together with Atg18^HA3-yEGFP^. Upon detergent lysis of whole cells and immunoadsorption on anti-mCherry beads, the three CSC subunits, but not the SNX-complex subunits, co-adsorbed Atg18^HA3-yeGFP^ (Fig. 1c). In cells lacking Vps26 (*vps26*Δ), the interaction between Atg18^HA3yeGFP^ and Vps35^yomCherry^ was decreased, yet it remained above the background defined by the non-tagged control (Fig. 1e). Thus, Vps26 contributes significantly but not exclusively to this interaction. In live cell confocal fluorescence imaging, Vps26^yomCherry^ and Atg18^yeGFP^ formed multiple puncta, which partially overlapped at the vacuolar membrane (Fig. 1d, white arrows). Triggering fragmentation of the vacuole through a moderate hypertonic shock increased the colocalization between Vps26 and Atg18, particularly at sites of vacuole-vacuole contact (Fig 1d). Both *in-vivo* and *in-vitro* observations thus suggest a novel complex between CSC and Atg18, which we term the CROP complex.

### Atg18 and sorting nexins compete for binding to Vps26/29/35

We tested the functional relevance of retromer for fission of vacuoles by labeling these organelles with the fluorophore FM4-64 in vivo. In agreement with previous observations (Raymond *et al*, 1992; Liu *et al*, 2012; Balderhaar *et al*, 2010), strains lacking the SNX subunits (*vps5*Δ and *vps17*Δ) presented many small vacuolar fragments whereas the CSC mutants *vps26*Δ, *vps29*Δ and *vps35*Δ showed fewer and larger vacuoles than the wild type (Fig. 2a). After addition of 0.5 M salt to stimulate vacuole fission (Bonangelino *et al*, 2002; Zieger & Mayer, 2012), all three CSC mutants maintained their large central vacuoles, whereas wildtype cells fragmented the compartment into multiple (>7) vesicles that were much smaller and numerous than before (Fig. 2b,c). This led us to the hypothesis that the CSC subunits promote vacuole fission whereas the SNX subunits may prevent hyper-activity of the fission machinery. They might do so by interfering with the formation of the CROP complex. In line with this, the co-immunoadsorption of Atg18^HA3-eGFP^ with Vps26^yomCherry^ was strongly increased in the SNX mutant *vps17*Δ (Fig. 3a). Furthermore, the vacuolar hyper-fragmentation of the *vps17*Δ cells could be fully reverted by deleting ATG18 and its redundant homolog ATG21, or by deleting VPS26 (Fig. 3 b,c). Thus, both CSC and Atg18 are required for vacuole fission.

**Figure 2:**
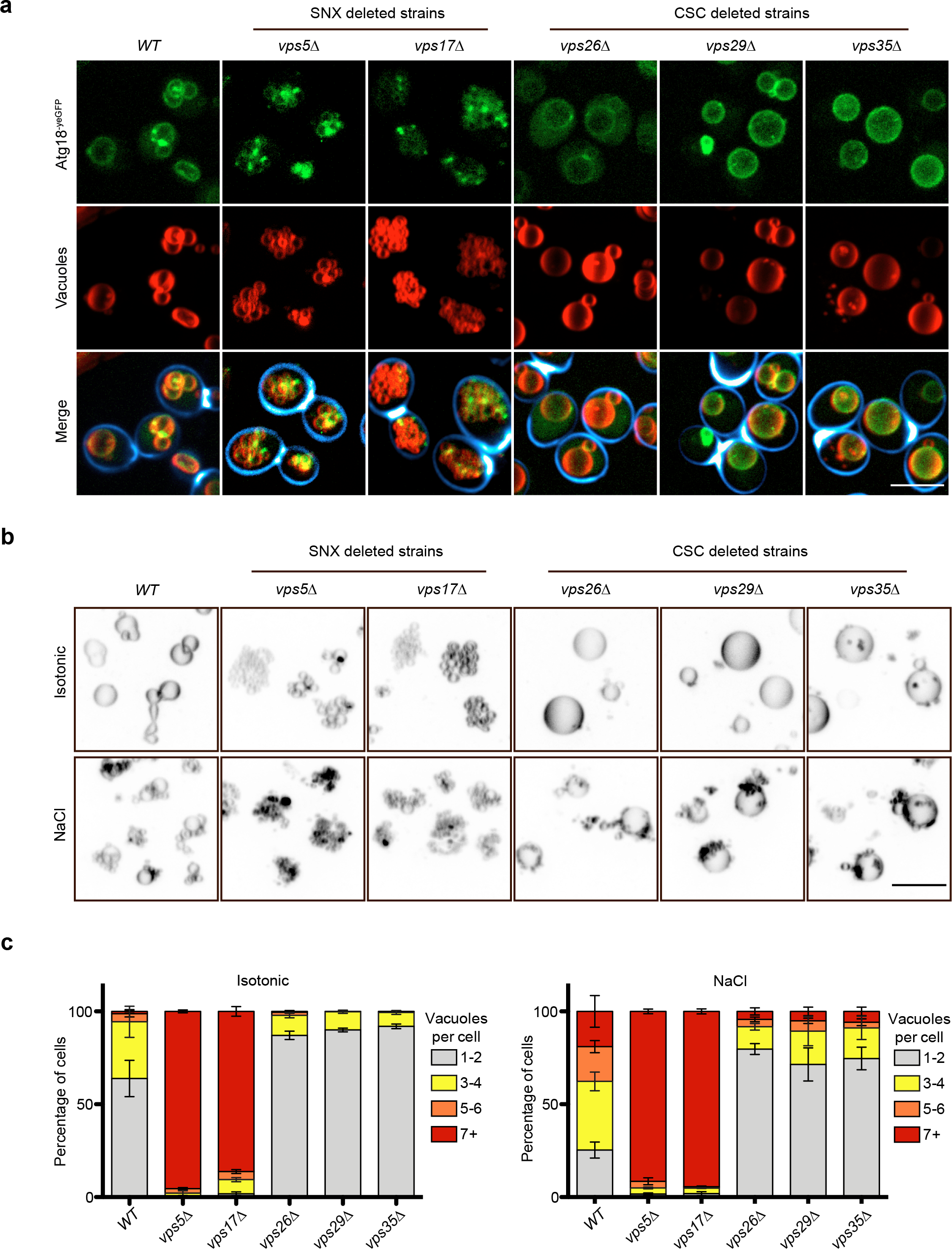
Effects of sorting nexins and CSC subunits on vacuole structure and vacuole fission *in vivo*. **a**, Vacuole structure. Cells carrying Atg18^yeGFP^ and the indicated retromer deletions were logarithmically grown in YPD medium, stained with FM4-64 and calcofluor white, and analyzed by confocal microscopy. Maximum intensity projections of z-stacks are shown. Scale bar: 5 µm. **b.** Salt-induced vacuole fission. The indicated cells were logarithmically grown in YPD and stained with FM4-64. Vacuole morphology was imaged as in a, before and after a mild salt shock with 0.5 M of NaCl for 15 min. The look-up table has been inverted to allow better representation of the clusters of extremely small vacuolar fragments in the SNX mutants. Scale bar: 5µm. **c**. The number of vacuoles per cell was quantified for the samples from b. n=3 experiments with at least 100 cells per condition were evaluated; Error bars represent the SEM.

**Fig. 3.**
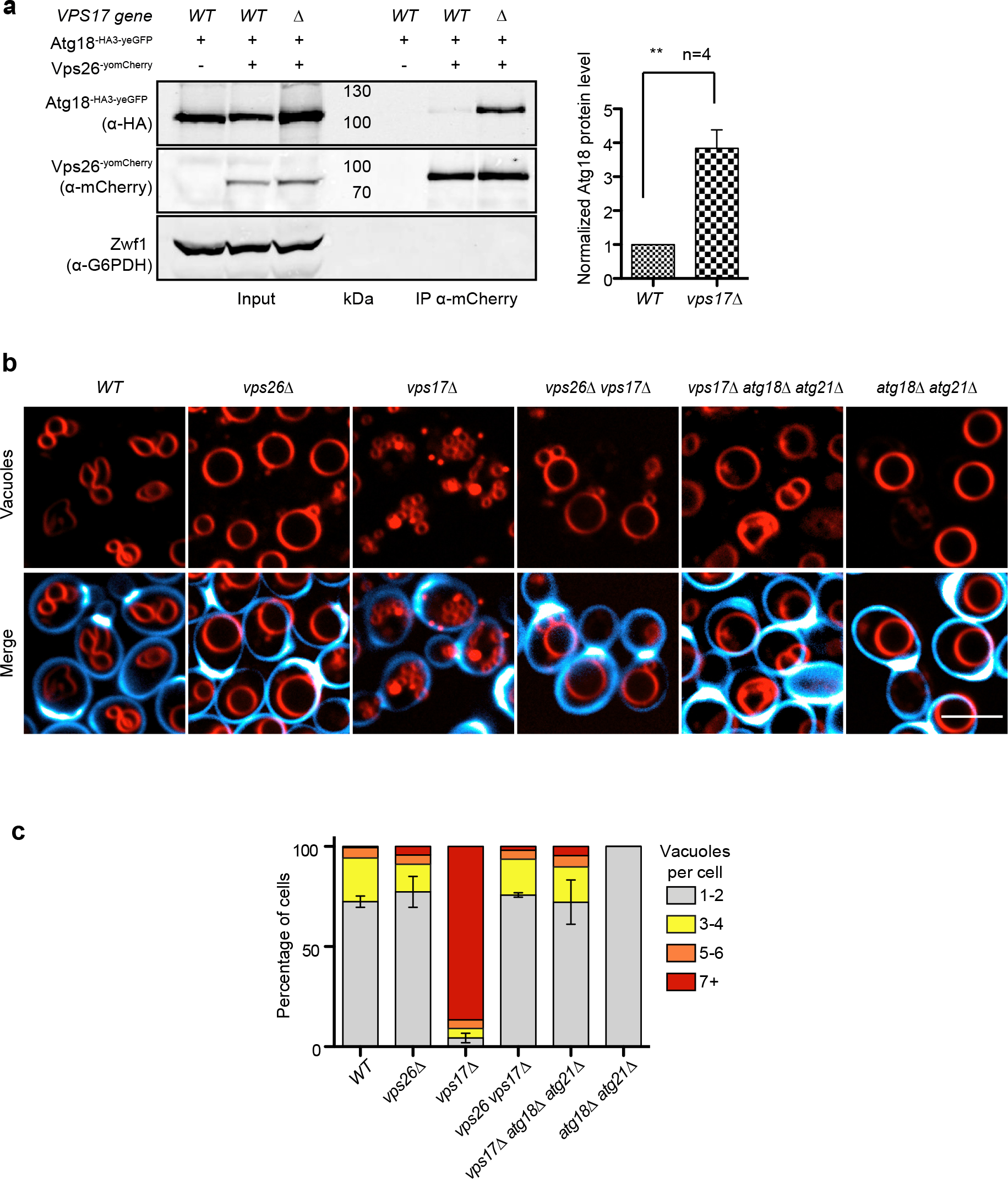
Interaction of Atg18 and CSC. **a**, Vps17 labilizes the Atg18-Vps26 interaction. Wildtype or *vps17*Δ cells expressing ATG18^HA3-yeGFP^ from a centromeric plasmid were logarithmically grown in YPD. Genomically tagged Vps26^yomCherry^ was pulled down from whole-cell extracts and analyzed for associated Atg18^HA3-yeGFP^ by SDS-PAGE and Western blotting. Glucose-6-phosphate dehydrogenase (Zwf1) serves as a loading control. The intensity of the interacting Atg18^HA3-yeGFP^ was quantified on a LICOR fluorescence imager and normalized to the amount of Vps26^yomCherry^. Means from n=4 independent experiments are shown; error bars represent the SEM. Values of the wildtype interaction were used as the reference and set to 1. **p<0.01 **b**, Epistasis of ATG18 and retromer genes concerning vacuolar morphology. The indicated cells were logarithmically grown in YPD medium, stained with FM4-64 and calcofluor white as in Fig. 2a and analyzed by confocal microscopy. Scale bar: 5 µm. **c**, Quantification of vacuole morphology. The number of vacuoles per cell was measured in the cells from b. The graph shows the fractions of cells displaying the indicated numbers of vacuolar vesicles. n=3 experiments with at least 100 cells per condition and experiment were analysed. Error bars represent the SEM.

To assay formation of CROP directly, we purified Atg18 from bacteria and CSC from yeast, where the Vps29 was labeled with GFP. We noted that the addition of Atg18 to CSC-GFP induced a strong shift in the fluorescence signal of the GFP. Ligand binding can change the properties of protein-bound fluorophores via changes in orientation, contacts, or rotational freedom of the fluorophore. Such changes are a common tool used in titration experiments to determine binding constants. Fluorescence intensity increased with recombinant Atg18 concentration, allowing us to estimate a K_d_ value close to 50 nM (Fig. 4b). We also evaluated the formation of CROP by blue native polyacrylamide gel electrophoresis and Western blotting. Here, Atg18 migrates mostly as expected for a monomer, but also shows a weaker band consistent with a dimer. Purified CSC forms two major bands, in line with previous observations of monomers and dimers (Kendall *et al*, 2020). Mixing CSC with an equimolar amount of Atg18 transformed CSC into more slowly migrating species, which both contained Atg18 and hence represent CROP (Fig. 4a). These bands were abolished, when purified SNX (Vps5/Vps17) was incorporated to the mix of proteins at a five-fold excess over Atg18 and CSC. This suggests that SNX interferes with CROP formation or stability, which is also consistent with our observation that CROP is more abundant in Atg18 pull-downs from *vps17*Δ strains (Fig. 3a).

**Figure 4:**
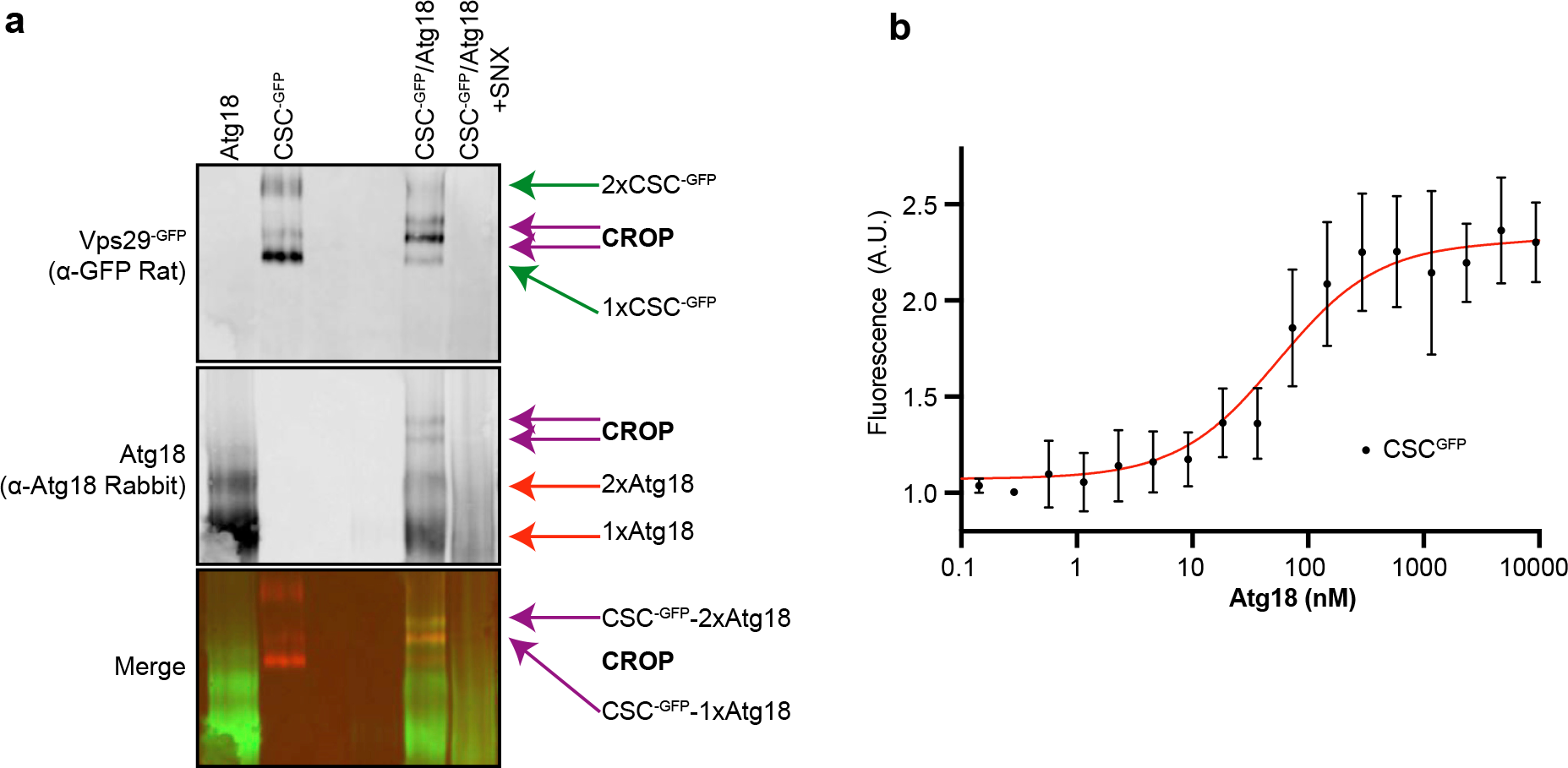
Formation of CROP from pure components and interference by SNX. **a**, Purified Atg18 and CSC^-GFP^ were mixed in a 1:1 ratio, in the presence or absence of a 5-fold excess of SNX. The proteins were incubated together in PBS and analyzed by native PAGE and Western blotting using the antibodies indicated in brackets. **b**, Pure recombinant Atg18 was titrated from 0 to 75 µM using the shift of fluorescence of Vps29^GFP^ (2.5 nM), which is induced by Atg18 binding. The curve was fitted using GraphPad Prism 9.

### The Atg18-CSC interaction is necessary for membrane fission

Our immuno-adsorptions occasionally yielded a slightly shorter, C-terminal proteolytic fragment of Atg18^HA3-yEGFP^. This truncated form had lost the capacity of full-length Atg18 to interact with CSC, suggesting that the binding site could be located in the removed N-terminal region, in blade 1 or 2 of the Atg18 ß-propeller. This region has also been implicated in binding Atg2, which is required for the function of Atg18 in autophagy (Watanabe *et al*, 2012; Rieter *et al*, 2013). Alignment of various Atg18 and WIPI1/2 orthologs revealed a stretch of conserved residues in blade 2 (Fig. 5a), which is located at the opposite side of the two phosphoinositide binding sites that anchor these proteins to the membrane (Fig. 5b). The motif contains three serines and threonines. At least two of these, Thr^56^ and Ser^57^, can be phosphorylated in vivo (Feng *et al*, 2015). We generated Atg18^HA3-yEGFP^ with alanine and glutamate substitutions of these residues and tested their consequences on vacuolar morphology and on the Atg18-Vps26 interaction (Fig. 5c). Except for S57E, which was hardly expressed, all other variants were expressed comparably as the wildtype, and they bound to vacuoles (Fig. 5d). A strong effect was observed for the T56E substitution. It largely abolished the co-immunoadsorption of Atg18^HA3-yEGFP^ and Vps26^yomCherry^, suggesting that it compromised the interaction of Atg18 with CSC. In vivo microscopy supported this: In contrast to Atg18^HA3-yEGFP^, which concentrates in numerous foci on the vacuole membrane, often at sites enriched in Vps26^yomCherry^, Atg18^T56E-HA3-yEGFP^ showed a homogenous distribution along the vacuole and no co-enrichment with Vps26^yomCherry^ (Fig. 5d). Thus, CSC is required to concentrate Atg18 in vivo.

**Figure 5:**
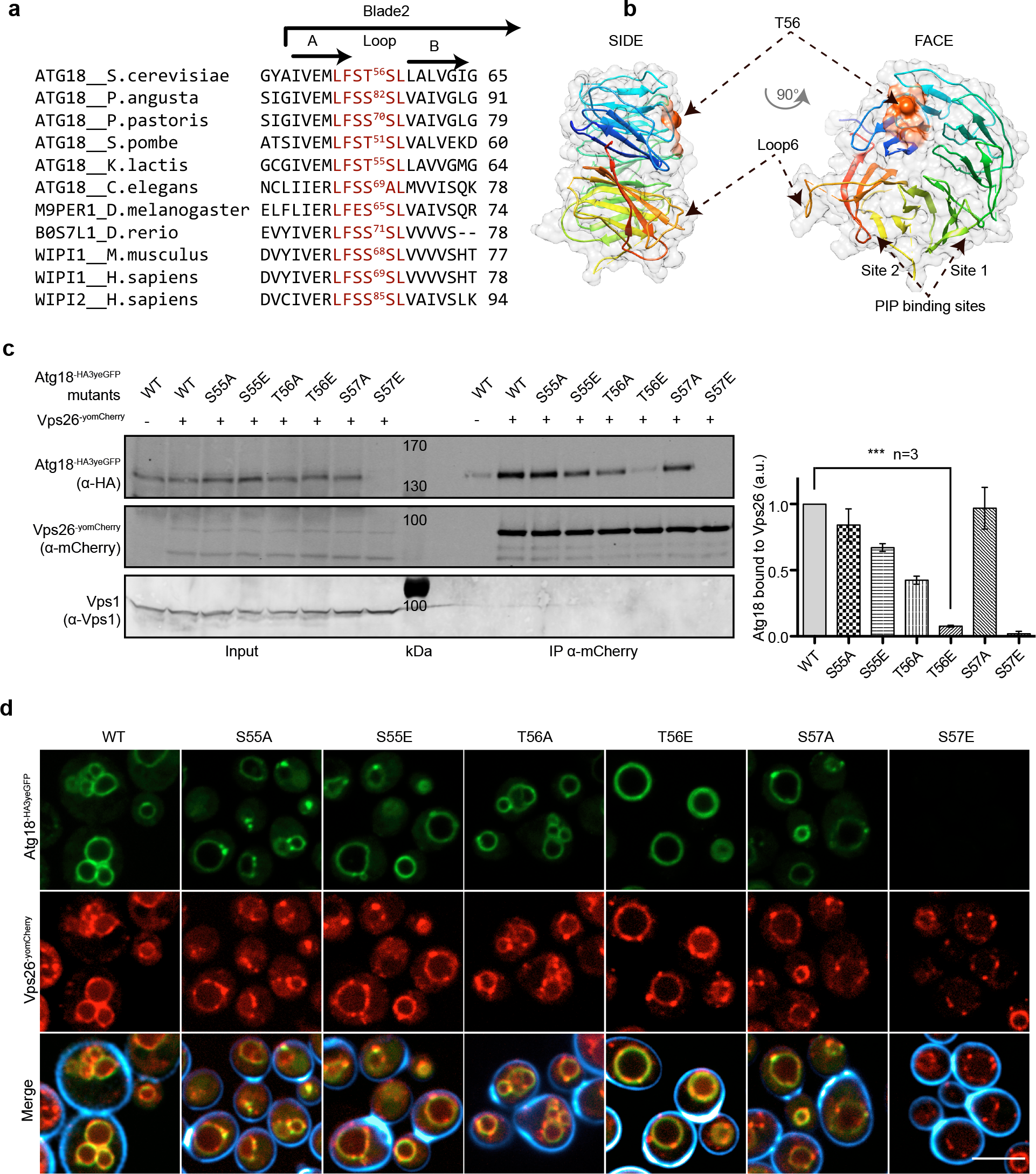
Substitutions labilizing the Atg18-Vps26 interaction. **a**, Sequence alignment of various Atg18 and WIPI1/2 orthologs showing a conserved stretch (in red) of residues in blade 2. **b**, The stretch containing T56 is mapped on the structure of Atg18 from *S. cerevisiae* (pdb #6KYB) (Lei *et al*, 2020), the LFSTSL motif in from *S. cerevisiae* is shown in orange, Thr56 in red. **c**, Pull-down. Cells (*SEY6210 atg18*Δ, *atg21*Δ) expressing genomically tagged Vps26^yomCherry^ and the indicated Atg18^HA3-yeGFP^ variants were logarithmically grown in SC^-URA^ media. Vps26^yomCherry^ was pulled down from whole-cell extracts with RFP-trap magnetic beads and analyzed by SDS-PAGE and Western blotting against the indicated proteins. Bands were quantified on a LICOR fluorescence imager. Signals of Atg18^HA3yeGFP^ were normalized relative to those of Vps26^yomCherry^. Vps1 served as a loading control. n=3 independent experiments were averaged. Error bars represent the SEM. *** p<0.001. **d**, Influence of the Atg18 substitutions on its localization. The cells from c were stained with calcofluor white to mark the cell walls and analyzed by confocal microscopy. The calcofluor signal (blue) is only shown in the merge. Scale bar: 5 µm.

The T56E substitution also impaired vacuole fission in vivo. Upon a salt shock, which stimulates rapid vacuole fission in *ATG18^WT^* cells, *atg18^T56E^* cells maintained few large vacuoles (Fig. 6 a,b). The hyper-fragmentation of vacuoles in *vps17*Δ cells, which depends on Atg18 (Fig. 3), provided an additional means of testing the effect of the T56E substitution. We used the *vps17*Δ *atg18*Δ *atg21*Δ cells, in which the hyper-fragmented phenotype of vps17Δ is suppressed by the lack of functional CROP. This allows the strain to recover 1-2 large vacuoles (see Fig. 3). Whereas re-expression of *ATG18^WT^* in this strain rescued vacuole fission and re-established hyper-fragmented vacuoles (Fig. 6 c,d), *atg18^T56E^* could not provide this activity and behaved similarly as the fission-defective *atg18^FGGG^*, a variant in which both phosphoinositide binding sites are compromised (Efe *et al*, 2007; Dove *et al*, 2004; Gopaldass *et al*, 2017; Baskaran *et al*, 2012; Krick *et al*, 2012) (Watanabe *et al*, 2012). These observations support the notion that the interaction of Atg18 with CSC in the CROP complex is necessary for vacuole fission.

**Figure 6:**
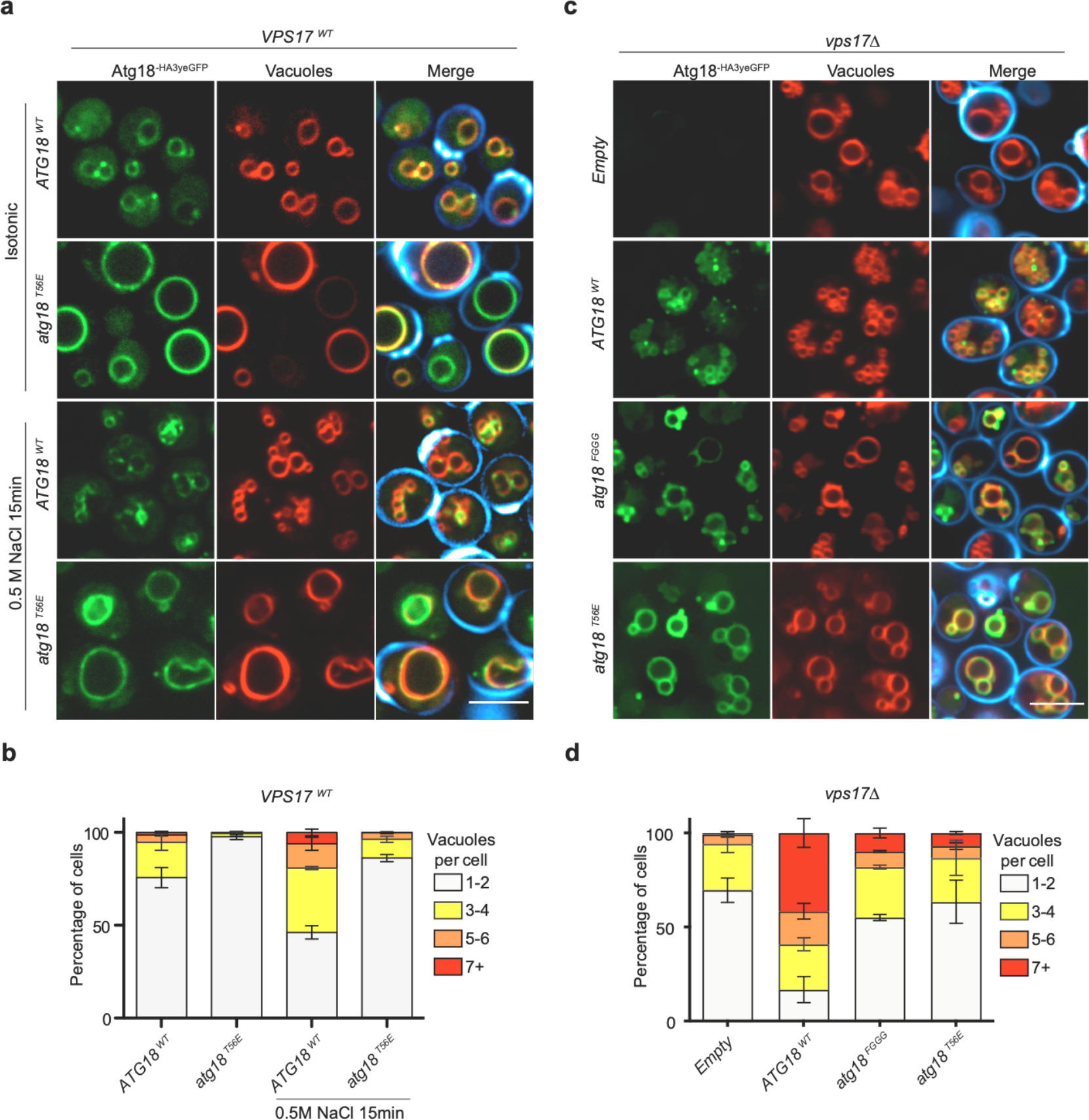
The Atg18-CSC interaction in CROP is essential for vacuole fission. **a**, Effect of Atg18^T56E^ on salt-induced vacuole fission.. Cells expressing Atg18^WTHA3-yeGFP^ or Atg18^T56E-HA3-yeGFP^ from centromeric plasmids in a *SEY6210 atg18*Δ, *atg21*Δ strain were logarithmically grown in SD^-URA^ medium. They were stained and imaged before and after the induction of vacuole fission by a short salt shock as in Fig. 1d. Calcofluor white-stained cell walls are only represented in the merge (blue). Scale bar: 5 µm. **b**, Quantification of the number of vacuoles per cell from a, n=3 experiments with at least 100 cells per condition were scored; error bars represent the SEM. **c**, Epistasis of *atg18^T56E^* over a *vps17*Δ mutation. The indicated variants of Atg18-HA3-yeGFP were expressed from plasmids in a *SEY6210 atg18*Δ, *atg21*Δ, *vps17*Δ strain. Cells were logarithmically grown in SD-URA and imaged as in Fig. 1d. Scale bar: 5 µm. **d**, Quantification of the experiments from c.

### CROP drives membrane fission on giant unilamellar liposomes

To directly test how CROP interacts with and acts on pure lipid membranes we created *in vitro* models with synthetic vesicles. Small unilamellar vesicles (SUVs) containing 5% each of PI3P and PI(3,5)P_2_ could recruit purified CROP components in a liposome centrifugation assay (Fig. 7a). Atg18^T56E^ and Atg18^WT^ fractionated with the liposomes whereas Atg18^FGGG^ bound very poorly. CSC by itself also interacted poorly with the vesicles. It was efficiently recruited to them through Atg18^WT^, but much less through Atg18^T56E^ and Atg18^FGGG^ (Fig. 7b). This provides evidence for the interaction of the pure CROP components on the membrane.

**Figure 7:**
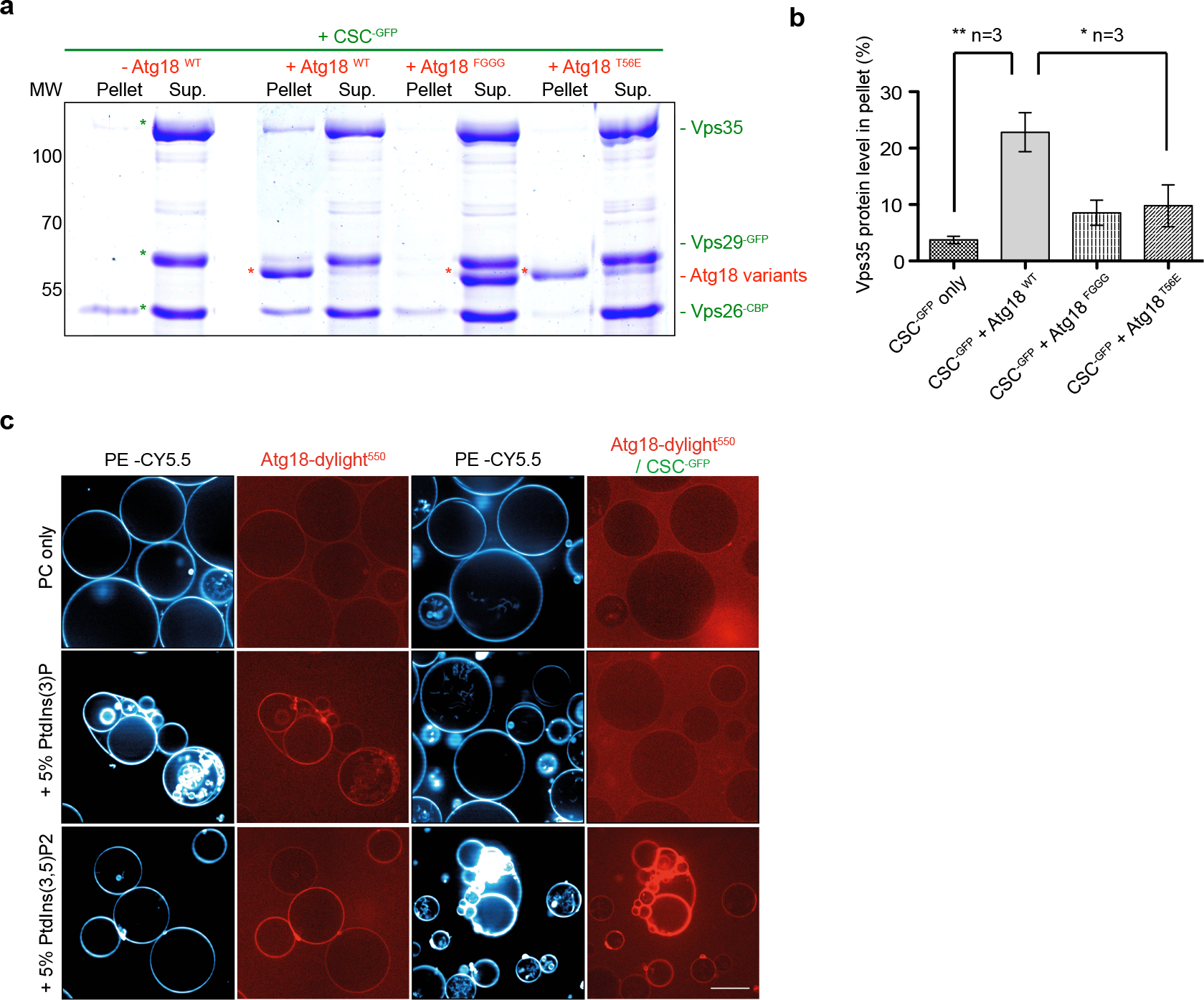
Impact of phosphoinositides on CROP binding. **a**, Atg18-dependent recruitment of CSC-GFP to small liposomes. SUVs were incubated (10 min, 25°C) with purified CSC (1.5 µM) alone or in combination with the indicated recombinant Atg18 variants (1.5 µM). The vesicles were sedimented by centrifugation and supernatants (Sup.) and pellets were analyzed by SDS–PAGE and Coomassie staining. **b**, Quantification by densitometry of the Coomassie signals on a LICOR scanner. ** p<0.001; * p<0.01 **c**, CROP recruitment to GUVs. GUVs containing the 5% of the indicated phosphoinositides and the fluorophore CY5.5-PE were left to sediment (30 min, 25°C) in wells that were supplemented with recombinant Atg18 (2.5 nM) covalently linked to dylight550 and CSC^GFP^ (100 nM) as indicated. Samples were incubated for 30 min before acquisition on a confocal microscope. Scale bar: 20 µm.

Next, we generated giant unilamellar vesicles (GUVs) from lipid mixtures containing 0.5% of the fluorescent lipid phosphatidylethanolamine-CY5.5. These vesicles are large enough for light-microscopic analyses (Fig. 7c). Size-fractionated GUVs were incubated with Atg18 that had been covalently coupled to a dylight^550^ fluorophore. Pure Atg18 rapidly bound the GUVs, but only when they contained PI3P or PI(3,5)P_2_, consistent with earlier studies showing that both lipids efficiently recruit PROPPINs to synthetic membranes (Baskaran *et al*, 2012; Krick *et al*, 2012; Scacioc *et al*, 2017; Gopaldass *et al*, 2017). Remarkably, Atg18 discriminated the two lipids when it was incubated in the presence of an excess of CSC, which incorporates Atg18 into CROP. Under these conditions, Atg18 bound only to the PI(3,5)P_2_-containing GUVs but not to PI3P-GUVs. This suggests that integration into CROP tunes the lipid affinity of Atg18 towards PI(3,5)P_2_, the phosphoinositide necessary to trigger vacuole fission in vivo.

Finally, we assessed the impact of CROP on GUV structure upon longer incubations. Atg18-dylight^550^ was bound to GUVs containing both 2.5% PI3P and 2.5% PI(3,5)P_2_. Thereby we sought to mimic the fact that the normally minor level of PI(3,5)P_2_ increases substantially when vacuole fission is induced by hypertonic shift. It can then become of similar abundance as PI3P (Bonangelino *et al*, 2002; Cooke *et al*, 1998). During the 30 min incubation, Atg18^WT^ and Atg18^T56E^ were recruited to the surface of the GUVs, whereas most Atg18^FGGG^ remained in the buffer (Fig. 8). Atg18^WT^ and Atg18^T56E^ distributed along the membrane in a homogeneous manner. Upon addition of 50 nM CSC-GFP, followed by a second incubation phase of 30 min, CSC bound to GUVs, when Atg18^WT^ was present. Binding coincided with the formation of a large number of small vesicles that remained attached to the GUVs. The generated vesicles showed strong signals of Atg18-dylight^550^ and CSC^GFP^, with CSC^GFP^ being concentrated in numerous puncta on or between these vesicles. Atg18^T56E^ recruited less CSC-GFP to the GUVs than Atg18^WT^, and it formed only very few smaller vesicles and CSC^GFP^ puncta. At the concentration used (25 nM), Atg18 alone did not induce any fission or tubulation on GUVs. Pure Atg18 can promote fission of GUVS only at >50 times higher concentration, as we have shown previously (Gopaldass *et al*, 2017). This suggests that the integration of Atg18 into CROP potentiates its membrane fission activity.

**Figure 8:**
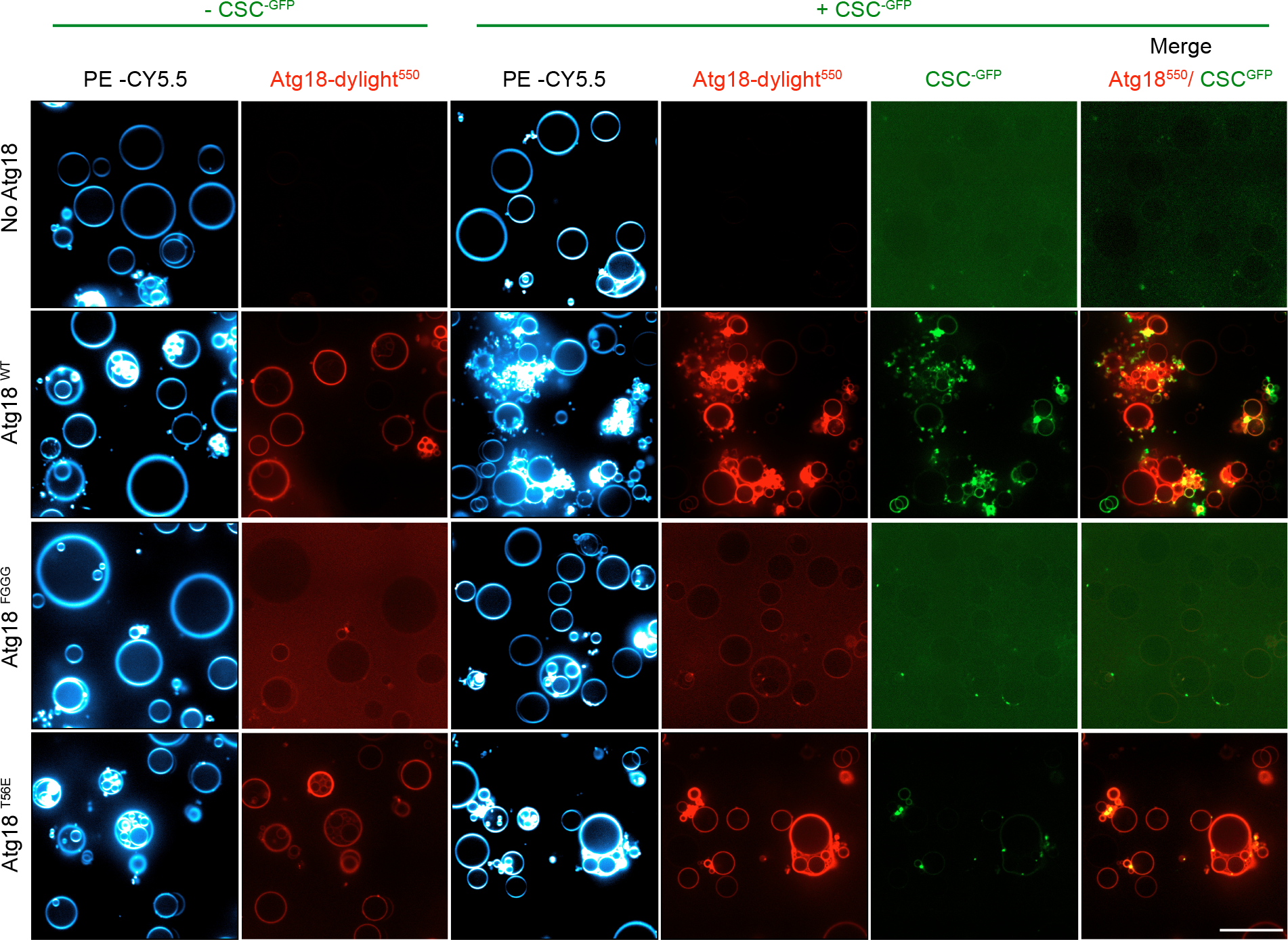
Fission of giant unilamellar liposomes by CROP. GUVs containing 2.5% PI3P, 2.5% PI(3,5)P_2_ and CY5.5-PE were incubated (30 min, 25°C) alone or with recombinant Atg18 variants (25 nM), which had been covalently linked to a dylight550 fluorophore (red). CSC^GFP^ (50 nM) was added to part of the samples for 30 min, before the vesicles were imaged on a confocal microscope. Scale bar: 20 µm.

### CROP is required for protein exit from mammalian endosomes

Both CSC and PROPPINs are conserved from yeast to mammals (in mammals, CSC alone is called retromer), suggesting that also CROP might be conserved. Sequence alignment and modeling allowed us to identify the equivalent residue of Atg18 T56 in its human homolog WIPI1 as the S69 residue (Fig. 5a). Since the endosomal system of mammalian cells is more developed than that of yeast (Day *et al*, 2018), it offers better possibilities to study the formation of endosomal transport carriers. This is illustrated by manipulations of WIPI1, which can suppress the fission of such carriers and lead to the accumulation of micrometer-long tubules on the endosomes of HK2 cells (De Leo *et al*, 2021). These tubules are very prominent and easy to recognize. We deleted WIPI1 from this human cell line and used plasmid transfection to re-express fluorescent WIPI1 fusions with S69A or S69E substitutions (Fig. 9a-c). Both variants were expressed to similar levels as the wildtype (Fig. 9c). While human Vps26^eGFP^ colocalized extensively with WIPI1^WT-mCherry^ in dots, which represent endosomes (De Leo *et al*, 2021), both WIPI1^S69^ substitutions partially segregated the two proteins, which is consistent with an impairment of their interaction. Similar observations were made using hVps35^eGFP^ instead of hVps26^eGFP^ (Supp. Fig 1a, b). In cells expressing WIPI1^S69A-eGFP^, and more so for those expressing WIPI1^S69E-eGFP^, micrometer-long membrane tubules emanated from endosomes. Their abundance and size were further increased by simultaneous knockdown of hVps35 (Fig. 9 d, e). Similar elongated tubules are observed when the fission activity of WIPI1 is abrogated by mutations in its lipid binding domains, or in its amphipathic helix in CD loop 6, which is essential for its fission activity (De Leo *et al*, 2021). Substitutions inactivating fission activity of WIPI1 (De Leo *et al*, 2021) also have effects on compartments carrying the lysosomal marker Lamp1. These were also recapitulated by WIPI1^S69E-eGFP^. Whereas lysosomes normally form small puncta that are dispersed in the cytosol and partially colocalize with eGFP-WIPI1^WT^, Lamp1-positive compartments are grossly enlarged in cells expressing WIPI1^S69E-eGFP^ (Supp. Fig 2b). eGFP-eWIPI1^S69A^ produced a qualitatively similar but weaker phenotype.

**Figure 9:**
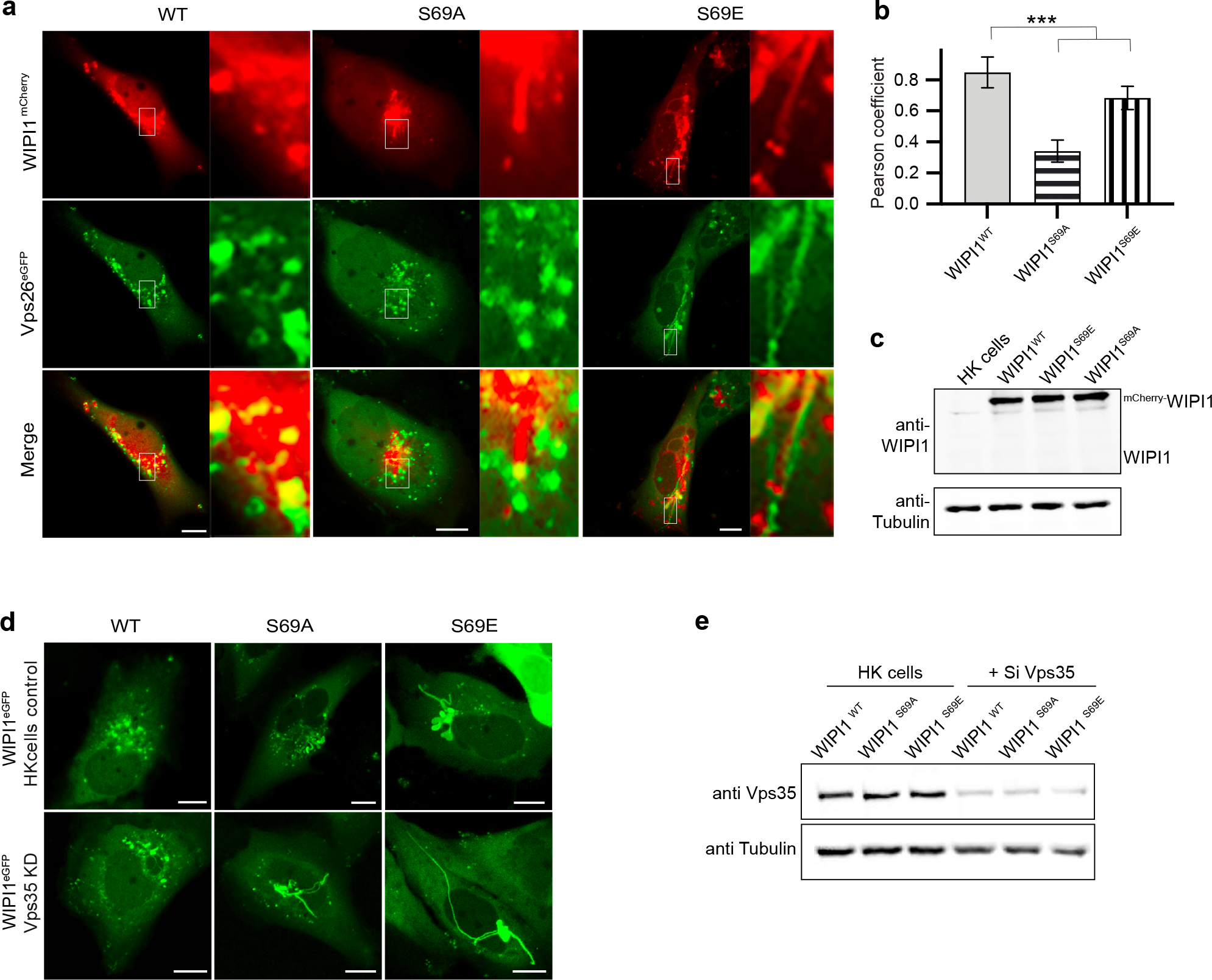
Effect of retromer and WIPI1 on human endosomes. **a**, Colocalization of WIPI1 with Vps26A. The indicated WIPI1^mCherry^ variants and Vps26^eGFP^ were expressed for 18 h in HK2 cells, from which endogenous WIPI1 had been deleted (WIPI1-KO). The cells were analyzed by confocal microscopy. Scale bars: 10 μm. Insets show enlargements of the outlined areas. **b,** Quantification of the colocalization in *a*, using the Pearson coefficient. Mean values ± SD are shown. n=3 independent experiments with a total of 150 cells were quantified per condition. *** p<0.01 **c,** Expression levels of WIPI1^mCherry^ variants. Lysates (50 μg of protein per sample) from the cells in A were analyzed by SDS-PAGE and Western blot against WIPI1 and tubulin. **d,** Additive effect of WIPI1^S69A^ and deletion of Vps35. HK2 cells expressing the indicated WIPI1^eGFP^ variants were transfected with siRNA against Vps35 or a control siRNA pool. Confocal microscopy was performed 18 h after transfection. Scale bar: 10 μm. **e**, Vps35 levels after knock-down. Lysates (50 μg per sample) from the cells in d were analyzed by SDS-PAGE and Western blot against Vps35 and tubulin.

We assayed the effects of the S69 substitutions on protein exit from the endosomes via trafficking of transferrin receptor (TfR), a protein that shuttles transferrin from the plasma membrane towards endosomes and back (Dautry-Varsat *et al*, 1983). The endosomes of HK2 cells were loaded with transferrin and then subjected to a chase in transferrin-free medium (Fig. 10 a,b). WIPI1 knockout cells re-expressing WIPI1^WT-eGFP^ efficiently returned transferrin back to the cell surface, from where it finally dissociates, leaving very little transferrin associated with the cells after the chase period. By contrast, cells expressing WIPI1^S69E-eGFP^ retained transferrin in their endosomes. WIPI1^S69E-eGFP^ was even dominant negative over the endogenous WIPI1, since it had a similarly strong effect on Tf recycling in wildtype cells (Supp. Fig. 2a) as in WIPI1 knockout cells (Fig. 10). That WIPI1^S69E-eGFP^ provokes the accumulation of exaggerated tubules and interferes with cargo exit from endosomes is consistent with the notion that CROP promotes fission of endosomal transport carriers at mammalian endosomes.

**Figure 10:**
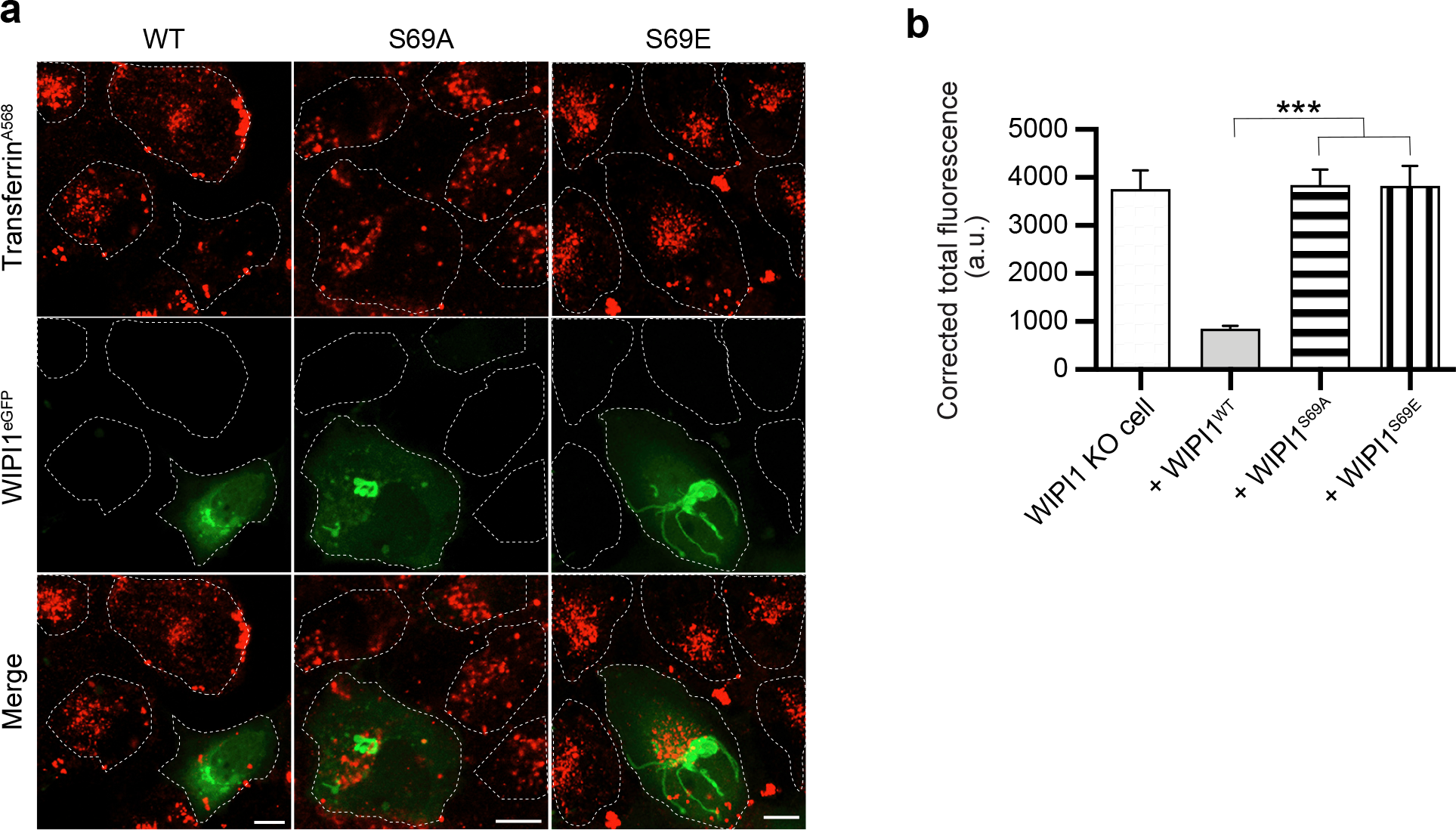
Effect of CROP on protein exit from mammalian endosomes. **a**, Tf recycling. WIPI1-KO cells were transfected with WIPI1^WT-eGFP^, WIPI1^S69E-eGFP^ or WIPI1^S69A-eGFP^ for 18h. Then, they were serum-starved for 1 h, loaded with Alexa Fluor 568-conjugated Tf, chased at 37°C for 1 h without labeled Tf, and analyzed by confocal microscopy. Scale bar: 10 μm. White dashed lines delineate the circumference of the cells. **b**, Quantification of Tf-fluorescence in the cells from a that expressed WIPI1 variants. Total cell fluorescence was integrated and corrected for background fluorescence. Mean values ± SD are shown. n=3 independent experiments with a total of 150 cells analyzed per condition. *** p<0.001.

## Discussion

Our results suggest that the PROPPIN Atg18 associates with parts of retromer to form the CROP complex. While pure Atg18 alone displays fission activity on GUVs at micromolar concentrations (Gopaldass *et al*, 2017), CROP promotes fission of these synthetic vesicles in the low nanomolar range, i.e. with much higher potency. In line with this, destabilization of CROP produces a number of in vivo phenotypes that are consistent with a loss of fission activity: It interferes with vacuole fragmentation and with endosomal membrane exit; and it leads to the accumulation of huge endosomal tubules, which were proposed to represent endosomal carriers that continue to elongate but fail to detach (De Leo *et al*, 2021). This favors the notion that CROP represents the relevant agent for fission on endo-lysosomal membranes in vivo. CROP provides a novel function to retromer subunits that are associated with Parkinson’s and Alzheimer’s disease, such as hVps35 (Li *et al*, 2019; McMillan *et al*, 2017; Rahman & Morrison, 2019). Therefore, it opens novel perspectives for the mechanistic analysis of these pathologies in relation to the fission activity of CROP.

The Vps26/29/35 complex (CSC) has a tendency to oligomerize (Hierro *et al*, 2007; Lucas *et al*, 2016; Kovtun *et al*, 2018; Kendall *et al*, 2020; Collins *et al*, 2008; 2005; Purushothaman *et al*, 2017). In the context of retromer, this oligomerisation supports the formation of tubular endosomal transport carriers, which sequester cargo exiting from endo-lysosomal compartments. If such oligomerisation occurred also when Atg18 is bound, it would concentrate multiple copies of Atg18 on a small membrane patch. This might be relevant for fission, because the hydrophobic CD loop 6 of Atg18 forms a conserved amphipathic alpha-helix when brought into contact with a bilayer (Gopaldass *et al*, 2017). The shallow insertion of this helix into the membrane should increase the curvature of the bilayer (Boucrot *et al*, 2012; Campelo *et al*, 2008). The concentration of several helices through oligomeric CSC is expected to enhance this effect, making fission more efficient.

Structural analyses of retromer and the associated SNX protein have yielded first models of how this coat might form tubular endosomal carriers (Hierro *et al*, 2007; Lucas *et al*, 2016; Kovtun *et al*, 2018; Kendall *et al*, 2020). The current models suggest a two-layered coat, in which an inner layer of SNX proteins recruits a peripheral layer of arch-shaped CSC complexes. Oligomerization of this coat is supported through multiple homo- and heteromeric interactions between CSC subunits, SNX subunits and cargo (Kovtun *et al*, 2018; Lucas *et al*, 2016) (van Weering *et al*, 2012). Our observations suggest that Atg18 and SNX compete for binding to CSC. If we assume that a retromer-coated tubule is a relatively homogeneous structure, in which all Vps35 and Vps26 subunits are engaged by SNXs, as proposed (Kovtun *et al*, 2018; Lucas *et al*, 2016), we can formulate a plausible working model for fission of ECVs. Since arch-like CSC structures carry Vps26 subunits at each of their “legs”, CSC might remain bound to the tubular coat that it has assembled through one of its legs, while on the other leg a PROPPIN could bind instead of a sorting nexin. Since the PROPPIN and sorting nexins appear to compete for binding, such a recruitment might be favored at the rim of the tubular SNX layer, i.e., at the site where fission should occur to detach an endosomal carrier. The competition for CSC binding with the SNXs might thus help to target CROP activity to the correct place.

Fission activity of CROP is not only used to facilitate departure of endosomal cargo, but it can also drive the division of an entire organelle, as shown by the fragmentation of vacuoles. In yeast, both reactions require not only CROP but also the dynamin-like GTPase Vps1 (Chi *et al*, 2014; Peters *et al*, 2004; Zieger & Mayer, 2012; Arlt *et al*, 2015). This is remarkable, because dynamin-like GTPases are mechanochemical devices, which can squeeze membrane tubules to very small radii (Antonny *et al*, 2016). We envision that the two protein systems cooperate to drive fission. This may create a situation similar to endocytosis, where the detachment of endocytic vesicles requires fission-promoting activity from dynamin and from additional membrane-deforming factors carrying fission activity, such as epsin (Boucrot *et al*, 2012). Dissecting the activities of CROP, retromer and dynamins will require refined in vitro systems that should allow to measure coat assembly, PROPPIN and dynamin recruitment, and membrane constriction.

## Materials and Methods

### Yeast cell culture

All strains were grown in either in YP (yeast extract, peptone) or in SC (synthetic dextrose) dropout media to select for auxotrophies and to avoid plasmids loss, both supplemented with 2% glucose. Conditions for SILAC growth are described below. Strains, plasmids and primers used in this study can be found in Table 2, 3 and 4. Liquid cultures were grown in at 30°C and shaken at 180 rpm.

**Table 1:** Atg18 interactors identified in the SILAC approach.

| Gene Name | Protein Name | Log2 Ratio (Standard media/Control) | Log2 Ratio (Hyperosmotic Shock/Control) | Log2 Ratio (Hyperosmotic shock/Standard media) |
| --- | --- | --- | --- | --- |
| *ATG18* | Autophagy-related protein 18 | 8.1 | 8.3 | 0.06 |
| *ATG2* | Autophagy-related protein 2 | 5.3 | 4.9 | -0.40 |
| *BUB1* | Checkpoint Serine/Threonine-protein kinase Bub1 | 4.9 | 4.5 | -0.34 |
| *SAP155* | SIT4-associated protein Sap155 | 3.5 | 4.2 | 0.56 |
| *GIS3* | Protein Gis3 | 1.6 | 2.7 | 0.97 |
| *SIT4* | Serine/threonine-protein phosphatase PP1-1 | 1.8 | 2.7 | 0.74 |
| *CDC55* | Protein phosphatase PP2A regulatory subunit B | 1.1 | 2.0 | 1.04 |
| *VPS35* | Vacuolar protein sorting-associated protein 35 | 1.1 | 1.6 | 0.60 |
| *VPS29* | Vacuolar protein sorting-associated protein 29 | 0.8 | 1.5 | 0.53 |
| *VPS26* | Carboxypeptidase Y-deficient protein 8 | 0.8 | 1.4 | 0.60 |
| *GIS2* | Zinc finger protein Gis2 | 0.5 | 1.2 | 0.68 |
| *ILV3* | Dihydroxy-acid dehydratase, mitochondrial | 0.5 | 1.0 | 0.45 |

**Table 2a.**
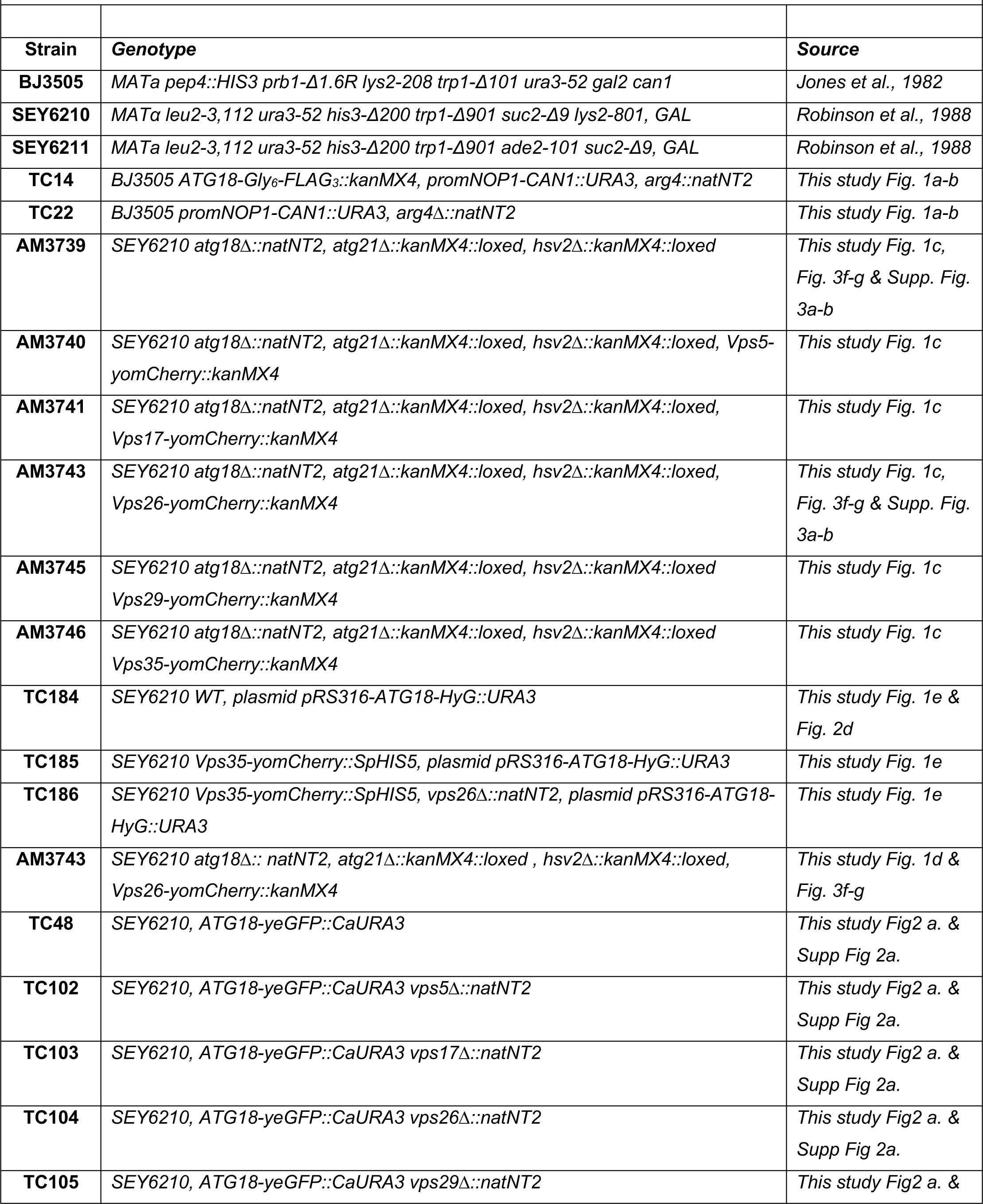
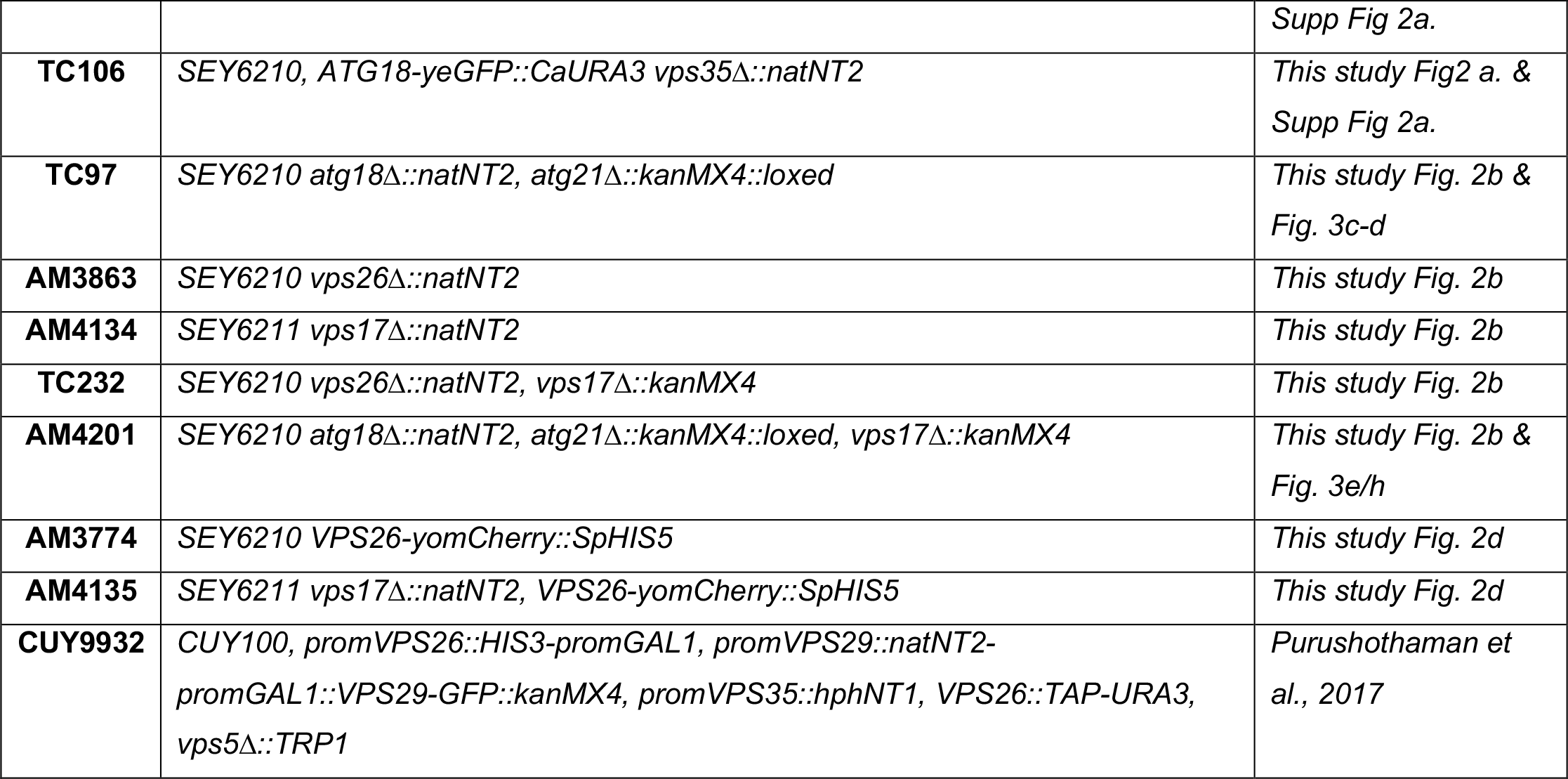
List of yeast strains.

**Table 2b.** List of mammalian cell lines.

| **Table 2b. List of mammalian cell lines** |  |  |
| --- | --- | --- |
| **MGDL** | HK-2 (ATCC® CRL-2190™) | *Fig. 5, Supp. Fig. 5 & 6* |
| **MGDL** | HK-2 , *wipi1*<sup>-/-</sup> | *This study Fig. 5 Supp. Fig. 5* |

**Table 3.**
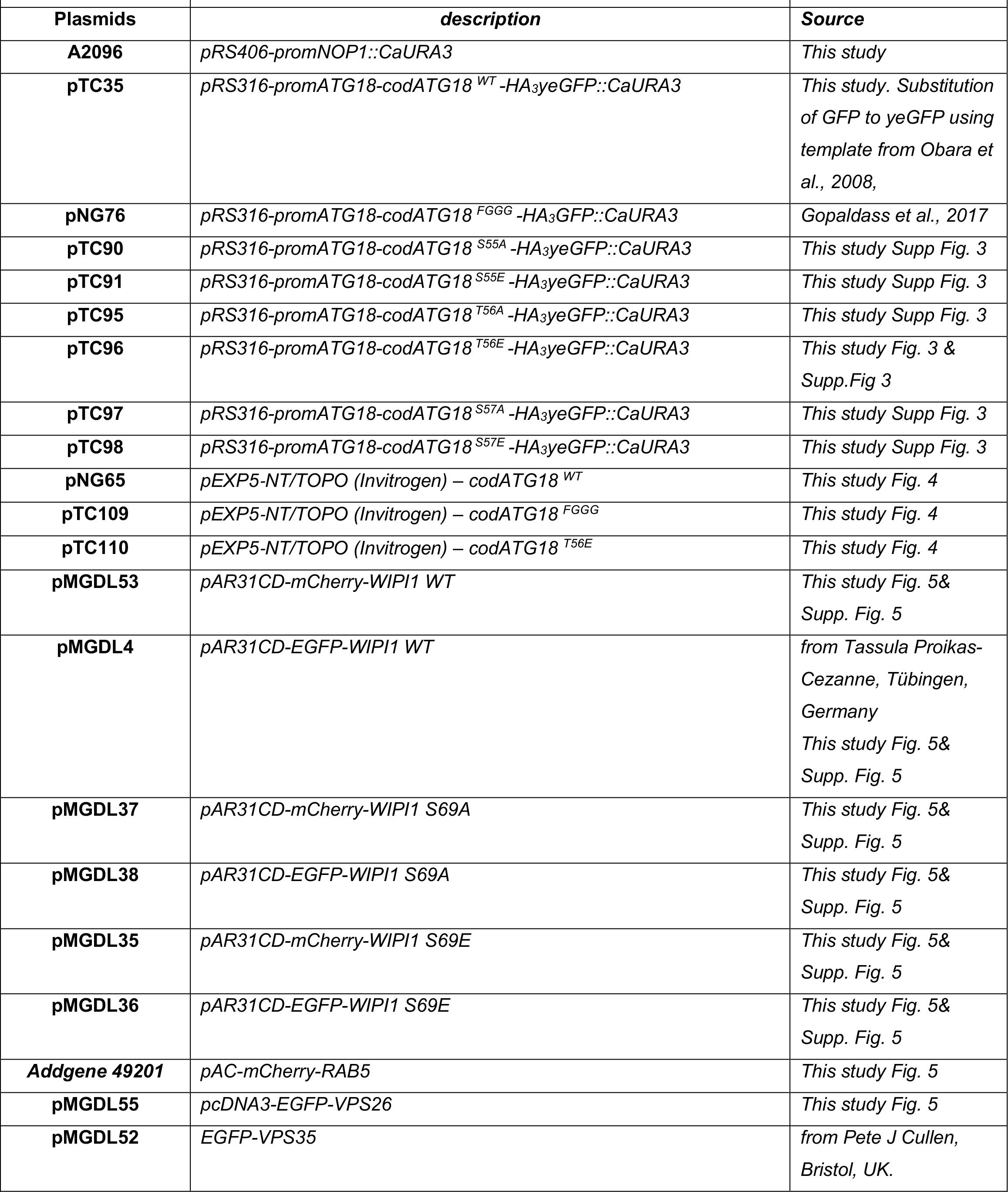
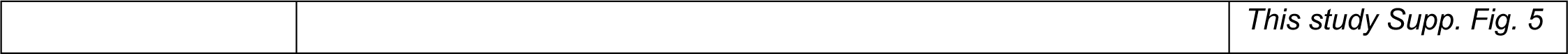
List of plasmids.

**Table 4.**
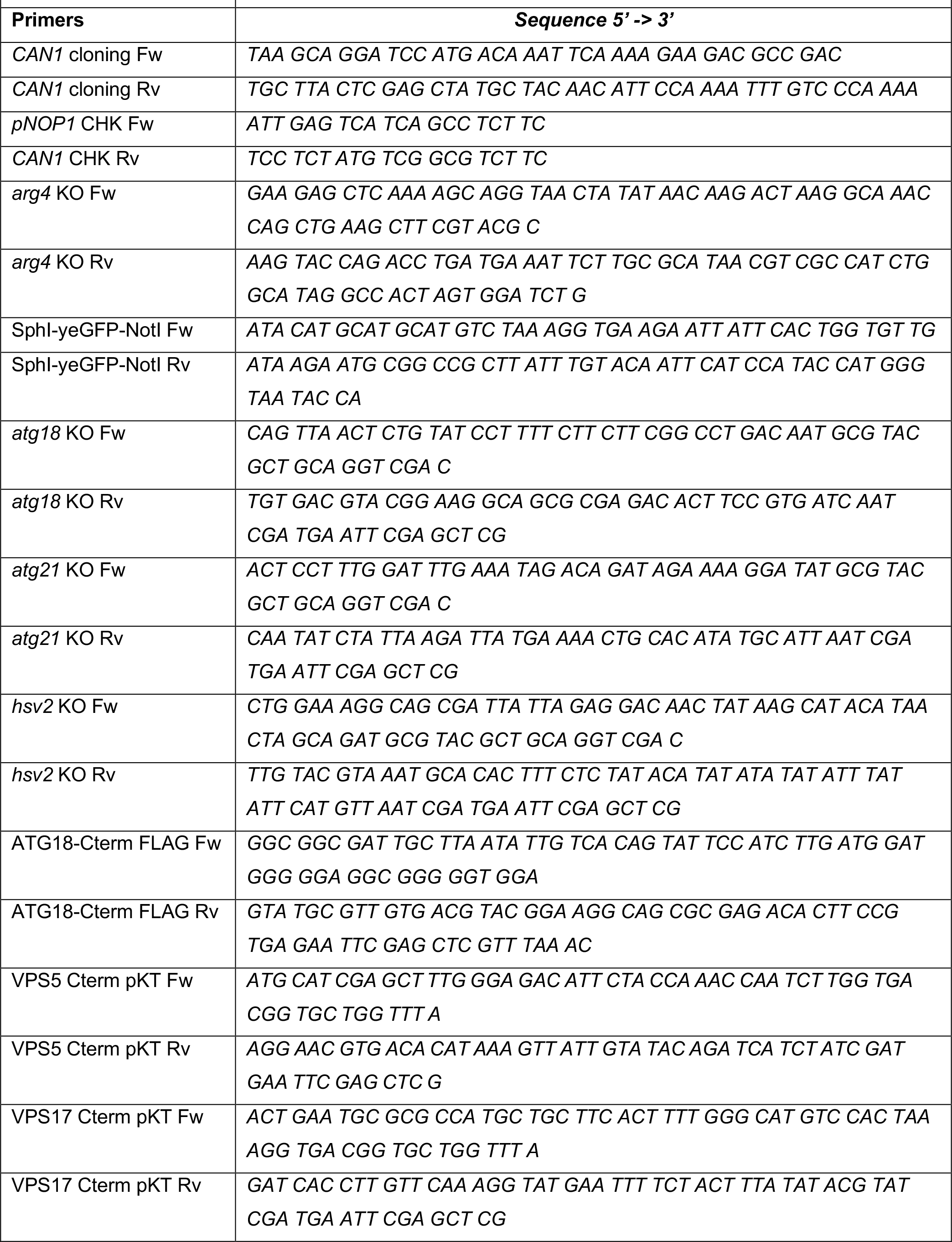
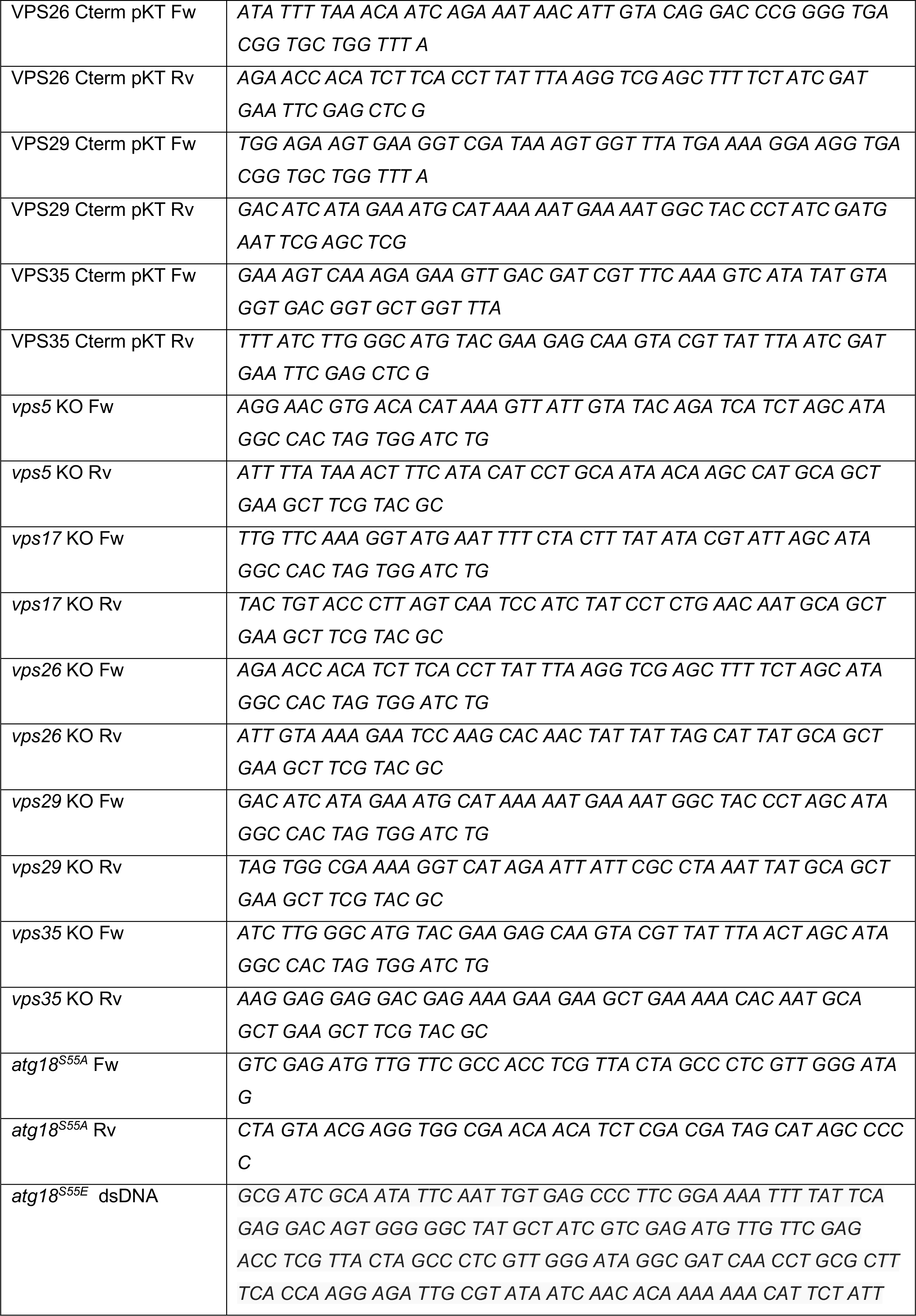
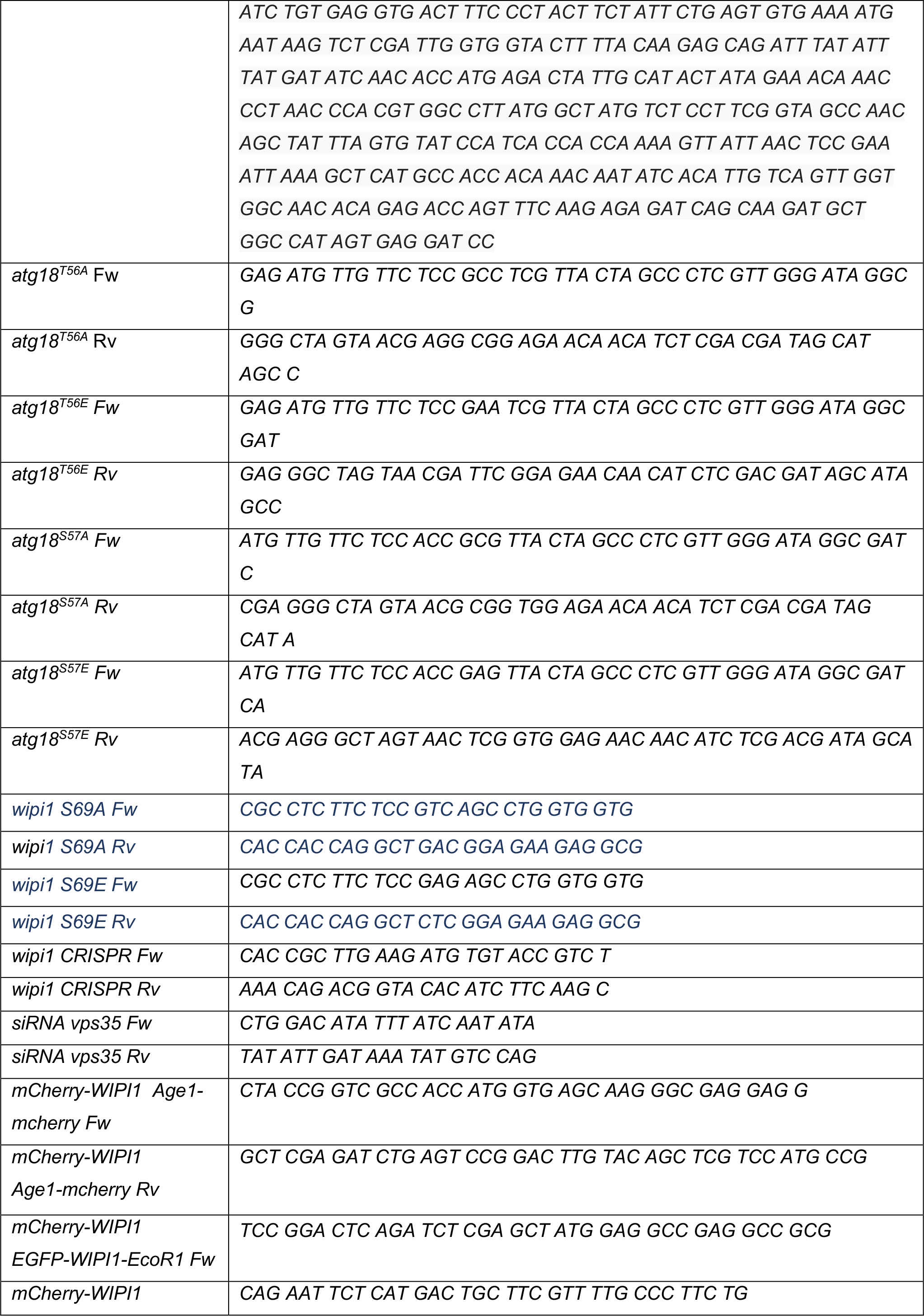
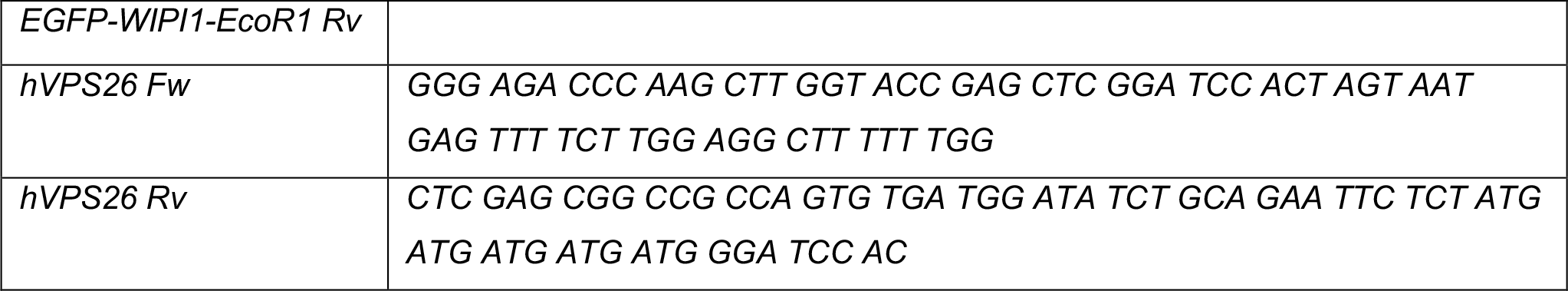
List of primers and double-stranded DNA.

### Strains and plasmids

Genes were deleted by replacing a complete open reading frame with a natNT2 (Euroscarf #P30346; Janke et al, 2004) or a loxP-flanked kanMX4 deletion cassette (Euroscarf #P30114; Güldener et al, 1996). Some constructs required to remove the kanMX4, using the Cre–lox P recombination method with plasmid pSH47 (Euroscarf #P30114; Güldener et al, 1996). Atg18 and CSC have been C-terminally tagged with either Gly6-FLAG3::kanMX4 (available on Addgene #20754; Funakoshi et al Yeast. 2009), yomCherry::kanMX4, and yomCherry::SpHIS5 (both available on Addgene #44903 & #44841; Lee et al., 2013) by direct insertion of these tags at their genomic locus. Atg18-GFP was expressed from pRS316-Atg18-HA3-GFP (Atg18-HG), which was a kind gift of Dr Y. Ohsumi (Tokyo Institute of Technology). GFP has been replaced for a yeast version yeGFP, subcloned from pKT0127 (Addgene #8728; Sheff et al, 2004) and placed in between the Sph1/Not1 restriction sites to generate pRS316-Atg18-HA3-yeGFP. Mutants in the putative retromer interaction motif (LFSTSL) of Atg18 were created by site-directed PCR mutagenesis (See Appendix for primer details), except for the mutant S55E, which has been generated by replacing the region between restriction site MfeI and MscI through chemically synthetized double-stranded DNA (from Eurofins) containing the mutation. Strains used for SILAC must be auxotrophic for lysine and arginine. BJ3505 already contains the lys2-208 auxotrophic and arginine auxotrophy was generated by deleting *ARG4*. Since BJ3505 cells lack activity of the arginine importer Can1, we restored uptake of exogenous arginine by expressing CAN1 from the NOP1 promoter. *CAN1* was subcloned from the BY4741 background into a *pRS406-promNOP1::CaURA3 plasmid*. This plasmid has been linearized using the StuI restriction site of the CaURA3 locus and transformed in BJ3505 *arg4*Δ, complementing the *ura3-52* mutation and creating the sTC22 strain suitable for SILAC. All constructs were verified by PCR and DNA sequencing; yeast transformations were performed using the LiAc/SS carrier DNA/PEG method (Gietz, R., Schiestl, R., *Nat Protoc* **2,** 31–34 (2007)

eGFP-WIPI1^WT^ (pAR31CD vector) was kindly provided by Tassula Proikas-Cezanne (Tübingen, Germany); EGFP-Vps35 was a gift from Peter Cullen (Bristol, UK). To generate mCherry-WIPI1, mCherry and WIPI1 fragments were amplified from pFA6a-mCherry-V5-KanMX6 (from Fulvio Reggiori, University Medical Center Groningen, Netherlands) and eGFP-WIPI1 plasmid, respectively, by using the primers Age1-mCherry/eGFP-WIPI1-EcoRI (see Table 4). The two fragments were fused by overlap extension-PCR and cloned into pAR31CD between Age1 and EcoR1 restriction sites.

eGFP-WIPI1^WT^ was used as DNA template for site-directed mutagenesis (QuikChange mutagenesis system, Agilent Technologies) to generate point mutations in the FSSS motif. mCherry-WIPI1^S69E^, mCherry-WIPI1^S69A^ were produced using mCherry-WIPI1^WT^ as template and eGFP-WIPI1^S69E^, and eGFP-WIPI1^S69A^ using eGFP-WIPI1^WT^ as template following the manufacturer’s protocol using primers (Microsynth) listed in Table 4. Non-mutated template vector was removed from the PCR mixture through digestion with Dpn1 for 1 h at 37°C. The product was purified by NucleoSpin PCR clean-up (Macherey-Nagel) and transformed into Escherichia coli. Plasmid DNA was purified and sequenced.

To generate Vps26-eGFP plasmids, we subcloned *hVPS26* from pmr101A-hVPS26 (Addgene #17636) with primer hVPS26 Fw / hVPS26 Rv (Table 4), into a pcDNA3-eGFP plasmid (Addgene #13031) using Gibson assembly.

### SILAC (stable isotope labelling by amino acids in cell culture)

Strains sTC14 and sTC22 were grown to saturation (approx. 1 day) in SC (synthetic complete, Formedium). 0.5 OD600 units were transferred into 5 mL of SC-arginine/-lysine (Sunrise Science Products) supplemented with 0.43 mM arginine and lysine. Light and stable isotope labeled amino acids (Sigma-Aldrich) were included in different conditions as described in the following table:

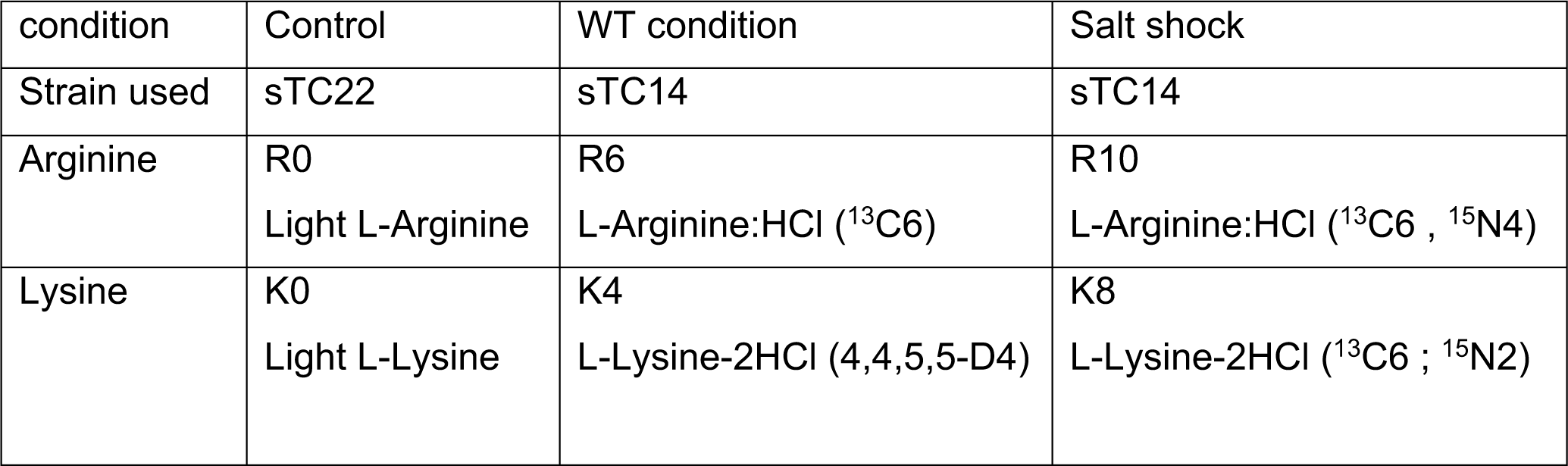

| condition | Control | WT condition | Salt shock |
| --- | --- | --- | --- |
| Strain used | sTC22 | sTC14 | sTC14 |
| Arginine | R0<br>Light L-Arginine | R6<br>L-Arginine:HCl (<sup>13</sup>C6) | R10<br>L-Arginine:HCl (<sup>13</sup>C6 , <sup>15</sup>N4) |
| Lysine | K0<br>Light L-Lysine | K4<br>L-Lysine-2HCl (4,4,5,5-D4) | K8<br>L-Lysine-2HCl (<sup>13</sup>C6 ; <sup>15</sup>N2) |

After 4 hours, these precultures were diluted into 1 Liter cultures. After 15 hours of culture at 30°C, vacuole fragmentation was triggered in the R10/K8 sample through a salt shock by addition of 200 mL of SC complete supplemented with 0.43 mM of R10/K8 and 5 M NaCl. After 5 minutes of shaking, cells were harvested with 5 min centrifugation (4800 × g, 4°C) in a Beckman JLA-8100 fixed-angle rotor. Cells were rinsed with ice-cold TGN buffer (50 mM Tris-Cl pH 7.4, 5% glycerol, 100 mM NaCl), and frozen in liquid nitrogen. Atg18 purification is described below. Before start of the experiments, tests were carried out to verify that medium and heavy samples were completely labeled (>99% labeling efficiency), and that no conversion of arginine to proline was observed.

### Protein purification

#### Atg18 purification for MS

For Atg18 purification in SILAC experiments, pellets were thawed on ice in one pellet volume of TGN lysis buffer (50 mM Tris pH 7.4, 10 % Glycerol, 100 mM NaCl) supplemented with 0.5% Triton, complete protease inhibitor tablets (Roche), phosphatase inhibitor tablets (Roche), 1 mM DTT, and 1 mM PMSF. Cells were passed one time through a French press (Constant Systems LTD) at 4°C with 2.2 kpsi of pressure. Atg18 was isolated by affinity purification using the Gly6-FLAG3 tag and Dynabeads (Sigma) crosslinked with the FLAG M2 antibody (F1804 epitope, Sigma). After 1 hour of incubation at 4°C, beads were washed 3 times with TGN buffer using a magnetic rack, transferred to new Eppendorf tubes, washed with 1 ml of elution buffer (Tris 50mM pH 7.4), then eluted with 3xFLAG peptide (0.5 mg/mL) in elution buffer. Eluates were flashed-frozen in liquid nitrogen and placed on a -80°C freezer before Coomassie staining and MS analysis.

#### Atg18 purification from bacteria

Atg18^WT^ and mutant DNA were amplified from the corresponding pRS316 plasmids and cloned into a pEXP5-NT/TOPO vector (Invitrogen). Plasmids were transformed into E. coli BL21. A 50 mL preculture overnight was used to inoculate 2 L of LB media (37°C). Cells were grown to an OD_600_ of 0.8–0.9. Cultures were then cooled to 16°C on ice, and IPTG (Roche) was added to a final concentration of 0.2 mM. Cells were shaken overnight (200 rpm, 16°C), pelleted, washed once in ice cold lysis buffer (500 mM NaCl, 50 mM Tris pH 7.4, 10 mM KPi), and resuspended in one pellet volume of lysis buffer with complete EDTA-free protease inhibitor cocktail (Roche) before purification. Purification was performed as previously described (Gopaldass *et al*, 2017). To conjugate the Dylight550 fluorophore (Thermofisher), proteins were incubated at room temperature equimolar with the dye in PBS containing 300mM of NaCl at room temperature protected from light. To remove the excess of fluorophore proteins were dyalised in PBS with 300mM NaCl overnight at 4°C with a 12kDa cutoff (ZelluTrans, ROTH).

#### Retromer purification from yeast cells

Strains for yeast purification were provided by Christian Ungermann (University of Osnabrück) (Purushothaman *et al*, 2017). Yeast precultures were grown in 50 mL YP-galactose (2%) to stationary phase, diluted in 2 L of YP-galactose and grown at least 24 h to late log phase (OD_600_ of 3). From this point, all steps were performed on ice or at 4°C. Cells were spun down (4800 x g, 5 min, 4°C) in a precooled Beckman JLA-8100 fixed-angle rotor, resuspended in one pellet volume of PBS (phosphate-buffered saline, pH 7.4) with 0.4 mM PMSF, pelleted as before, frozen in liquid nitrogen and stored at -80°C. Pellets were thawed in one pellet volume of RP buffer (50 mM Tris pH 8.0, 300 mM NaCl, 1 mM MgCl_2_, 1 mM PMSF, Roche Complete protease inhibitor tablet 1x). Cells were opened using a French press (Constant Systems LTD, pressure 2.2 kpsi). 5 mg DNase I (from Bovine Pancreas grade II, Roche) was added to 50 mL of lysate. The lysate was incubated on rotating wheel at 4°C for 20 min, and pre-cleared by centrifugation at 18’500 × g for 30 min in a JLA 25.50 rotor. The supernatant was cleared by centrifugation at 177’520 × g for 90 min in a Beckmann Ti60 rotor. After aspiration of the upper lipid phase, the cleared supernatant was passed through a 0.22 µM filter (Millipore) and transferred to a new 50 mL falcon tube. Cleared lysate was incubated with 1 mL IgG Sepharose beads suspension (6 Fast Flow, GE Healthcare) pre-rinsed with buffer for 20 ml lysate on a rotating wheel at 4°C for 1 h. Beads were spun down at 3000 × g for 5 min with minimal deceleration (Eppendorf 5804R) and washed 3 times with RP buffer, then transferred to new 1.5 mL Eppendorf tube. Beads were resuspended in 1mL of RP buffer without inhibitors, supplemented with 250 µg of TEV protease. TEV cleavage was performed at 16°C for 1h. The supernatant was incubated for an additional 20 min at 16°C with Ni-NTA beads to remove TEV protease. The supernatant was then concentrated on an Amicon Ultra-50 100K filter column at 3000 × g (Eppendorf 5804R) to reach a final volume of 250 µL. Eluates were divided into 10 µl aliquots, frozen in liquid nitrogen and stored at -80°C until use.

### MS analysis and MS data analysis

#### Sample preparation

The light-, medium- (Arg6/Lys4) and heavy-labeled (Arg10/Lys8) samples were mixed and concentrated by a microspin column with a 10 kDa cutoff membrane. 2/3 (for shotgun analysis) and 1/3 (for ubiquitination analysis of atg18 protein) of the mixed sample was fractionated in separate lanes of a 12% SDS-PAGE gel. For shotgun analysis, the whole lane was cut in 7 bands and digested, as described (Shevchenko *et al*, 1996) with sequencing-grade trypsin (Promega). For the ubiquitination analysis, only two bands between 50 and 150 kDa were cut and digested, using methyl methanethiosulfonate (MMTS) for cysteine alkylation.

#### Mass spectrometry analyses

After digestion, the extracted peptides were analyzed either on a hybrid LTQ Orbitrap Velos (for the shotgun analysis) or an Orbitrap Fusion Tribrid (for the ubiquitination analysis) mass spectrometer (Thermo Fisher Scientific, Bremen, Germany) connected to an Ultimate 3000 RSLC nano HPLC system (Dionex, Sunnyvale, CA, USA). Solvents used for the mobile phase were 97:3 H2O:acetonitrile (v/v) with 0.1 % formic acid (A) and 20:80 H2O:acetonitrile (v/v) with 0.1 % formic acid (B). Peptides were loaded onto a trapping microcolumn Acclaim PepMap100 C18 (20 mm x 100 μm ID, 5 μm, 100Å, Thermo Scientific), before separation on a reversed-phase custom packed nanocolumn (75 μm ID × 40 cm, 1.8 μm particles, Reprosil Pur, Dr. Maisch). A flowrate of 0.3 μl/min was used with a gradient from 4 to 76% acetonitrile in 0.1% formic acid, with a total method time of 95 (Velos) or 65 min (Fusion). In the Velos instrument, full survey scans were performed at a 60’000 resolution (at m/z 400). In data-dependent acquisition controlled by Xcalibur software (Thermo Fisher Scientific), the 10 most intense precursor ions detected in the full MS survey performed in the Orbitrap (range 350-1500 m/z) were selected and fragmented. MS/MS was triggered by a minimum signal threshold of 3’000 counts, carried out at relative collision energy of 35 % and with isolation width of 4.0 amu. Only precursors with a charge higher than one were selected for CID fragmentation and fragment ions were analyzed in the ion trap. The m/z of fragmented precursors was then dynamically excluded from any selection during 30 s.

In Fusion mass spectrometer, full survey scans (range 350-1550 m/z) were performed at a 120’000 resolution (at m/z 200), and a top speed precursor selection strategy was applied by Xcalibur software to maximize acquisition of peptide tandem MS spectra with a maximum cycle time of 3.0s. MS/MS was triggered by a minimum signal threshold of 5’000 counts. HCD fragmentation mode was used at a normalized collision energy of 32%, with a precursor isolation window of 1.6 m/z. Only precursors with a charge between 2 and 6 were selected for fragmentation and MS/MS spectra were acquired in the ion trap. Peptides selected for MS/MS were excluded from further fragmentation during 60s.

#### Data analysis

The SILAC triplex data were processed by the MaxQuant software (version 1.5.1.2) (Cox & Mann, 2008) incorporating the Andromeda search engine (Cox *et al*, 2011)). A UniProt yeast (Saccharomyces cerevisiae, strain ATCC 204508 / S288c) proteome database was used (downloaded in October 2014, 6’674 sequences), supplemented with sequences of common contaminants. Trypsin (cleavage at K,R) was used as the enzyme definition, allowing 2 missed cleavages.

For Velos data, carbamidomethylation of cysteine was specified as a fixed modification. N-terminal acetylation of protein, oxidation of methionine and phosphorylation of serine, threonine and tyrosine were specified as variable modifications. For Fusion data, methylthiolation of cysteine was specified as a fixed modification. N-terminal acetylation of protein, oxidation of methionine and ubiquitination of lysine were specified as variable modifications. All identifications were filtered at 1% FDR at both the peptide and protein levels with default MaxQuant parameters. MaxQuant data were further processed with Perseus software (Tyanova *et al*, 2016). Raw data is available in Appendix 2.

### Retromer co-immunoadsorption

To pull down retromer, 50 mL cultures in SC-URA with 2% D-Glucose were inoculated from a stationary 24-hour pre-culture in the same medium in and grown overnight to logarithmic phase (OD_600_ nm of 0.5 to 1). Cells were spun in 50 ml falcon tubes in a pre-cooled centrifuge (3000 × g / 5 min / 4°C). From this step, all manipulations were performed on ice or in the cold room. Pellets were resuspended in 1 ml ice-cold TGN lysis buffer (50 mM Tris pH 7.4, 10 % glycerol, 100 mM NaCl) supplemented with 0.5% Triton X-100, containing complete protease inhibitor tablets 1x (Roche), phosphatase inhibitor tablets 1x (Roche), 1 mM DTT, and 1 mM PMSF, and transferred to 2 ml Eppendorf tubes (round bottom). Cells were pelleted on a bench-top centrifuge at 800 x g for 3 min (Microfuge 16 – Beckmann Coulter). After discarding supernatants, cells were resuspended with 200 µl of lysis buffer and 100 μl of acid-washed glass beads (Sigma–Aldrich). Tubes were vigorously shaken for 10 min on an IKA Vibrax shaker (IKA, Staufen, Germany). After a quick spin, 200 ul of cold TGN lysis buffer was added to the lysate and the lysate was removed from glass beads using a 200 ul pipette tips to avoid transferring glass beads to the new 1.5 ml Eppendorf tube. The lysate was spun for 10 min at 10’000 × g on a cold bench-top centrifuge. Clear lysates were carefully recovered, without touching pellets, and transferred to new 1.5 ml Eppendorf tubes. Protein concentration was assayed in a Nanodrop1000 (Thermo Fisher Scientific) at 280 nm. Protein concentrations were equilibrated between samples by diluting with cold TGN lysis buffer, resulting in 500 µl of lysates (∼10 mg protein), of which 30 µl were kept as “Input” controls, and 470 µl was incubated with 20 µl pre-rinsed (with lysis buffer) RFP-Trap magnetic beads (Chromotek). After a 1-hour incubation on a rotating wheel, beads were pelleted using a magnetic separation rack. Beads were washed 3 times with TGN lysis buffer and transferred to a new Eppendorf tube. After discarding the supernatant, 20 ul of deionized water and 20 µl of NuPAGE 4X (Thermo-Fisher) supplemented with 100 mM of DTT were added to the beads. Inputs and eluates were denatured at 90°C for 10 min. After denaturing, beads were pelleted with a magnetic rack and supernatants were transferred to a new Eppendorf tube to form the “IP” samples. Inputs and IP were loaded on 10% SDS-polyacrylamide gels.

### SDS–PAGE and Western blotting

Lysates and eluates from Immunoprecipitations were run on 10% acrylamide gels for SDS-PAGE, freshly prepared and used the same day: 10% Protogel (30% w/v acrylamide, 0.8% bisacrylamide (37.5:1 solution, National diagnostics, Atlanta, USA), 380 mM Tris–HCl pH 8.8, 0.1% w/v SDS (Applichem, Darmstadt, Germany), 0.06% v/v TEMED (Applichem), 0.06% w/v APS (Applichem) for the running gel and 5% Protogel, 165 mM Tris–HCl pH 6.8, 0.1% w/v SDS, 0.08% v/v TEMED, 0.04% w/v APS for the stacking gel. Running buffer for SDS–PAGE was 190 mM glycine (Applichem), 25 mM Tris-base (Applichem), 0.5% SDS. To facilitate Atg18 migration and avoid formation of aggregates, samples were reduced and denatured at 90°C using NuPAGE buffer (Thermofisher) containing LDS instead of SDS and supplemented with 100 mM DTT. Gels were blotted on 0.45 µm nitrocellulose membrane (Amersham) overnight at a constant current of 200 mA using a Trans-Blot® Cell (Bio-Rad, USA). Membranes were decorated using anti-mCherry-1C51 (Abcam), anti-HA.11-16B12 (BioLegend), anti-G6PDH (Sigma-Aldrich), anti-Tubulin (clone B5-1-2, Sigma-Aldrich) and anti-WIPI1 (C-terminal epitope, Sigma-Aldrich).

### Native gel electrophoresis

For native gel electrophoresis, purified proteins were mixed, incubated at 25°C for 30 min, supplemented with loading buffer (50 mM BisTris pH 7.2, 6 N HCl, 50 mM NaCl, 10% w/v glycerol) and incubated for further 10 min. Then, samples were loaded on precast Bis-Tris polyacrylamide 4-16% gradient gels (Thermo Fisher Scientific). Electrophoresis buffers contained 50 mM BisTris, 50 mM Tricine pH 6.8 and were applied to anode buffer reservoirs. Cathode reservoirs were supplemented in addition with 0.002% Coomassie G-250. Electrophoresis was carried out at 4 °C for 90 min at a constant voltage of 150 V. Then, voltage was increased to 250 V for 60 min. Gels were Western blotted overnight with constant current of 200 mA using a Trans-Blot® Cell (Bio-Rad, USA). Membranes were destained from residual Coomassie traces by washes in methanol for several minutes, before blocking and antibody decoration.

### Atg18-CSC dissociation constant

The dissociation constant of CROP was determined by measuring the fluorescence intensity of a GFP coupled to Vps29. We noted that the fluorescence properties of this tagged protein changed because of Atg18 binding and exploited this effect to follow the binding event. Pure recombinant Atg18 was titrated from 0 to 75 µM by serial 1:1 dilution in PBS and supplemented with 2.5 nM of CSC-GFP, giving a final volume of 100 µL. After incubation at 30°C during 30 min, fluorescence was measured in microtiter plates using a SpectraMax Gemini EM spectrofluorometer (excitation 488 nm, emission 525 nm, cutoff 520 nm). Experiments were repeated three times. K_d_ was determined using nonlinear regression curve fitting (One site-Total) in GraphPad Prism9.

### Live microscopy and vacuole fragmentation

#### Live-cell imaging, FM4-64 staining, and fragmentation assay

Cells were inoculated from a stationary pre-culture (SC-URA or YP) supplemented with 2% D-glucose and grown overnight to logarithmic phase (OD_600_ nm between 0.5 and 1). After dilution to OD_600_ = 0.5 in 1 ml culture, 10 μM FM4-64 was added from a 10 mM stock in DMSO. Cells were incubated (1 h, 30°C, 180 rpm), followed by three washing steps with medium without FM4-64 (2 min, 3,000 × g) and a chase of 1h in medium without FM4-64. For induction of vacuole fragmentation, NaCl was added to the media to a final concentration of 0.4 M and cells were imaged at 0, 15, and 30 min after its addition. Cells were removed from the shaker, concentrated by a brief low-speed centrifugation, placed on a glass microscopy slide and imaged immediately. Z-stacks with a spacing of 0.3 µm were recorded on a NIKON Ti2E Yokogawa spinning disc confocal microscope with a 100x 1.49 NA objective and Photometrics Prime BSI cameras. Image analysis was performed with ImageJ.

#### Liposome preparation and microscope imaging

Lipids were purchased from Avanti Polar Lipids (USA): Egg L-a-phosphatidylcholine (EggPC); 1,2-dioleoyl-sn-glycero-3-phospho-L-serine sodium salt (PS); 1,2-dioleoyl-sn-glycero-3-phospho-(1′-myo-inositol-3′-phosphate) (PI3P); 1,2-dioleoyl-sn-glycero-3-phospho-(1′-myo-inositol-3′,5′-bisphosphate) (P(3,5)P_2_, 1,2-dioleoyl-sn-glycero-3-phosphoethanolamine-N-(Cyanine 5.5) (Cy5.5-PE) and porcine brain polar lipid extract (PL). All lipids were dissolved in chloroform and phosphatidylinositol phosphates were dissolved in chloroform/methanol/water (20:10:1).

Small unilamellar vesicles (SUVs) contained phospholipids in the following ratios: 89.5% PL + 5% PI3P + 5% PI(3,5)P_2_ supplemented with 0.5% Cy5.5-PE. SUVs were prepared as described (Baskaran *et al*, 2012). Lipids from stock solutions were diluted in chloroform in a glass tube and the solvent evaporated under argon flux while gently vortexing the tube. Tubes were dried at 55°C in vacuum for 1 h in order to remove traces of solvents. Retromer SUV buffer (20 mM HEPES pH 6.8, 50 mM KAc, 130 mM Sucrose, 10 µM ZnCl_2_) was added (final lipid concentration 5 mg/ml), and the tubes were placed in an oven at 37°C to hydrate the lipid film for 1 hour. After vortexing, lipids were transferred to an Eppendorf tube and frozen and thawed three times using liquid nitrogen. SUVs were prepared on the day of experiment or kept at −20°C and used within a week. SUVs were incubated with purified CSC-GFP (1.5 µM) for 10 min at room temperature (25°C) alone or in addition with recombinant Atg18^WT^, Atg18^FGGG^ or Atg18^T56E^ (1.5 µM). The suspension was spun for 10 min at 10’000 × g in a bench-top centrifuge and the supernatants and pellets were analyzed by SDS–PAGE and Coomassie staining.

Giant Unilamellar Vesicles (GUVs) were made with the following lipid composition: 89.5% PC, 5% PS, 2.5% PI3P, 2.5% PI(3,5)P_2_), supplemented with 0.5% Cy5.5-PE. To prepare GUVs, we followed the electro-formation method (Angelova et al, 1992). Freshly prepared lipid mix in chloroform was deposited on indium-titan oxide glass slides and dried for 1 hour at 55°C to evaporate all solvents in a vacuum oven. After mounting a chamber from 2 glass slides and an O-ring filled with a 250 mM saccharose solution, GUVs were electro-formed at 1 V and 10 Hz for 60 min at 55°C.

GUV solution was removed from the chamber by careful pipetting with a cut tip and placed in a 1.5 ml siliconized microcentrifuge tube. To purify the GUVs and remove excess lipids, GUVs from two chambers were pooled and an equivalent volume of retromer GUV buffer (20 mM HEPES pH 6.8, 115 mM KAc, 10 µM ZnCl_2_) was added to facilitate sedimentation in an Eppendorf swing bucket rotor for microcentrifuges (200 × g / 5min / RT). The supernatant was removed without touching the GUV pellet (∼50 ul). After washing a second time with the retromer GUV buffer, the supernatant was removed, and GUVs were resuspended in 150 µl of retromer GUV buffer. GUVs were used immediately for imaging.

For imaging, 10 μl of GUV suspension was added to 50 μl of retromer GUV buffer containing Atg18 protein at 25 nM concentration in a 96-well clear bottom plate (Greiner Bio-One, Thermo Fisher Scientific) pre-coated with solution of BSA dissolved in water (1 mg/ml) for 30 min. GUVs were left to sediment for 30 min. Imaging was done with a NIKON Ti2E spinning disc confocal microscope. Large image acquisition was generated by automatically stitching 5x5 fields from multiple adjacent frames. A minimum of 5 acquisitions were performed in different regions of the well. Picture analysis was performed with ImageJ.

### Mammalian cell experiments

All chemical reagents were from Sigma-Aldrich unless otherwise specified. Other reagents: Opti-MEM and Trypsin (Gibco® by Life Technologies); Alexa Fluor®568-conjugated Transferrin from Human Serum (Thermo Fisher Scientific).

#### Cell culture, transfection and treatments

HK2 cells were grown in DMEM-HAM’s F12 (GIBCO-Life Technologies); supplemented with 5% fetal calf serum, 50 IU/mL penicillin, 50 mg/mL streptomycin, 5 μg/mL insulin, 5 μg/mL transferrin, 5 ng/mL selenium (LuBio Science). Cells were grown at 37°C in 5% CO_2_ and at 98% humidity. Media, serum and reagents for tissue culture were purchased from GIBCO (Invitrogen).

HK2 cells were transfected with different plasmids using X-tremeGENE HP DNA transfection reagent (Sigma-Aldrich) according to the manufacturer’s instructions and incubated for 18-24 h before fixation or live-cell imaging. The HK-2 cell line was checked for mycoplasma contamination by a PCR-based method. All cell-based experiments were repeated at least three times.

#### Knockouts and RNA interference

HK2 WIPI1-KO cells were produced by using the CRISPR/Cas9 system as described (De Leo *et al*, 2021). For RNA interference, HK2 cells were transfected with siRNA for 72 h using Lipofectamine RNAiMax (Thermo Fisher Scientific) according to the manufacturer’s instructions. The siRNA targeting VPS35 was from Sigma (5’ CTGGACATATTTATCAATATA 3’; 3’ TATATTGATAAATATGTCCAG 5’). It was used at 20 nM final concentration. Control cells were treated with identical concentrations of siGENOME Control Pool Non-Targeting from Dharmacon (D-001206-13-05).

#### Transferrin recycling

HK2 cells were serum-starved for 1 h at 37 °C, washed twice in cold PBS with 1% BSA, and then exposed to 100 μg/ml Alexa-Fluor-488-Tf for 1 h at 37 °C (LOAD). After extensive washing with complete fresh HEPES-buffered serum free-medium, the recycling of Tf was followed by incubating the cells in Tf-free complete medium (CHASE) for 1 h at 37 °C. The cells were acid-washed (150 mM NaCl, 10 mM acetic acid, pH 3.5) before fixation.

### Immunostaining

HK2 cells were grown to 70% confluence on glass coverslips before immunofluorescence microscopy was performed. Cells were fixed for 8 min in 0.2% glutaraldehyde and 2% paraformaldehyde in PBS. This condition favors preservation of tubular endosomal structures although not to the extent seen by live microscopy. After fixation, cells were permeabilized in 0.1% (w:v) saponin (Sigma-Aldrich, 558255), 0.5% (w:v) BSA and 50 mM NH4Cl in PBS (blocking buffer) for 30 min at room temperature. The cells were incubated for 1 h with primary antibodies (anti-LAMP1 H4A3 from USBiologicvak Life Sciences) in blocking buffer, washed three times in PBS, incubated for 1 h with the secondary antibody (Cy3-conjugated AffiniPure Donkey anti-Mouse IgG H+L from Jackson Immuno Research), washed three times in PBS, mounted with Mowiol (Sigma-Aldrich, 475904-M) on slides and analyzed by confocal microscopy.

#### Confocal fluorescence microscopy and image processing

Confocal microscopy was performed on an inverted confocal laser microscope (Zeiss LSM 880 with airyscan) with a 63x 1.4 NA oil immersion lens. Z-stack Images were acquired on a Zeiss LSM880 microscope with Airyscan. Tf-fluorescence was quantified in CTR and WIPI1-KO cells using ImageJ. z-stacks were compressed into a single plane using the ‘maximum intensity Z-projection’ function in Image J. Individual cells were selected using the freeform drawing tool to create a ROI (ROI). The ‘Measure’ function provided the area, the mean grey value and integrated intensity of the ROI. The mean background level was obtained by measuring the intensity in three different regions outside the cells, dividing them by the area of the regions measured, and averaging the values obtained. This background noise was removed from each cell, yielding the CTCF (corrected total cell fluorescence): CTCF=integrated intensity of cell ROI − (area of ROI × mean fluorescence of background).

To quantify the degree of co-localisation, confocal z-stacks were acquired, single channels from each image in 8-bit format were thresholded to subtract background and then the “Just Another Colocalisation Plug-in” (JACOP) of ImageJ was used to measure the Pearson’s correlation coefficient.

## Statistics

Where averages were calculated, the values stemmed from experiments that were performed independently. Significance of differences was tested by a two-tailed t-test.

## Acknowledgements

We thank Véronique Comte-Misérez for assistance in protein purification, Manfredo Quadroni for the MS analysis of the SILAC experiment, and C. Ungermann for strains expressing CSC and SNX.

## Author contributions

TC performed Atg18 interaction studies and yeast experiments. Liposome experiments were performed by TC. Their estabishment and analysis was supported by NG. MGDL performed experiments with mammalian cells. AM conceived the study. All authors analyzed data and jointly wrote the manuscript.

---

**Supplementary Figure 1:**
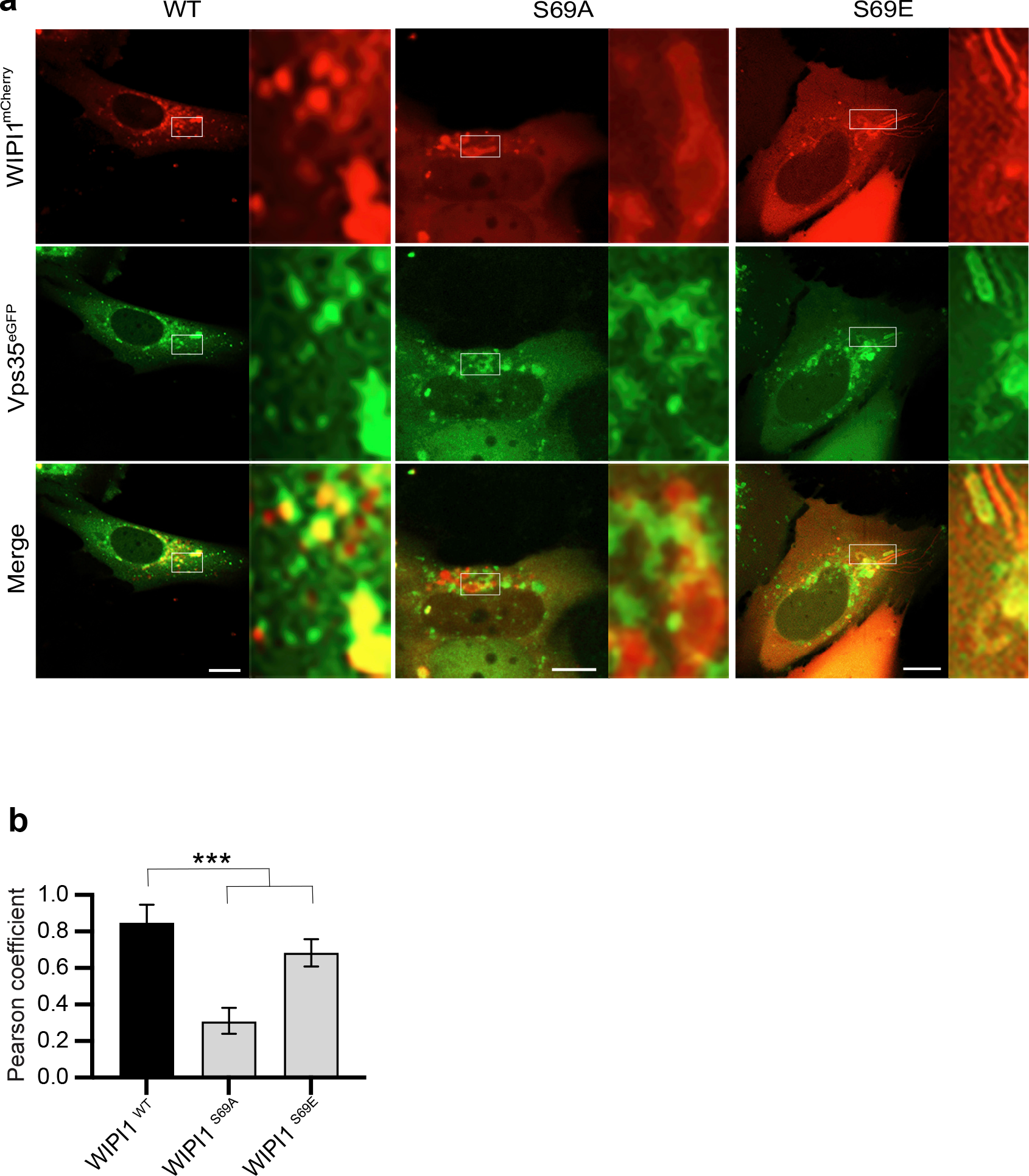
Colocalization of WIPI1^S69^ variants with hVps35. **a**, The indicated WIPI1^S69mCherry^ variants and Vps35^eGFP^ were expressed for 18 h in HK2 cells, from which endogenous WIPI1 had been deleted. The cells were analyzed by confocal microscopy. Scale bars: 10 μm. Insets show enlargements of the outlined areas. **b,** Quantification of the colocalization in *a*, using the Pearson coefficient. Mean values ± SD are shown. n=3 independent experiments with a total of 195 cells were quantified per condition. *** p<0.001.

**Supplementary Figure 2:**
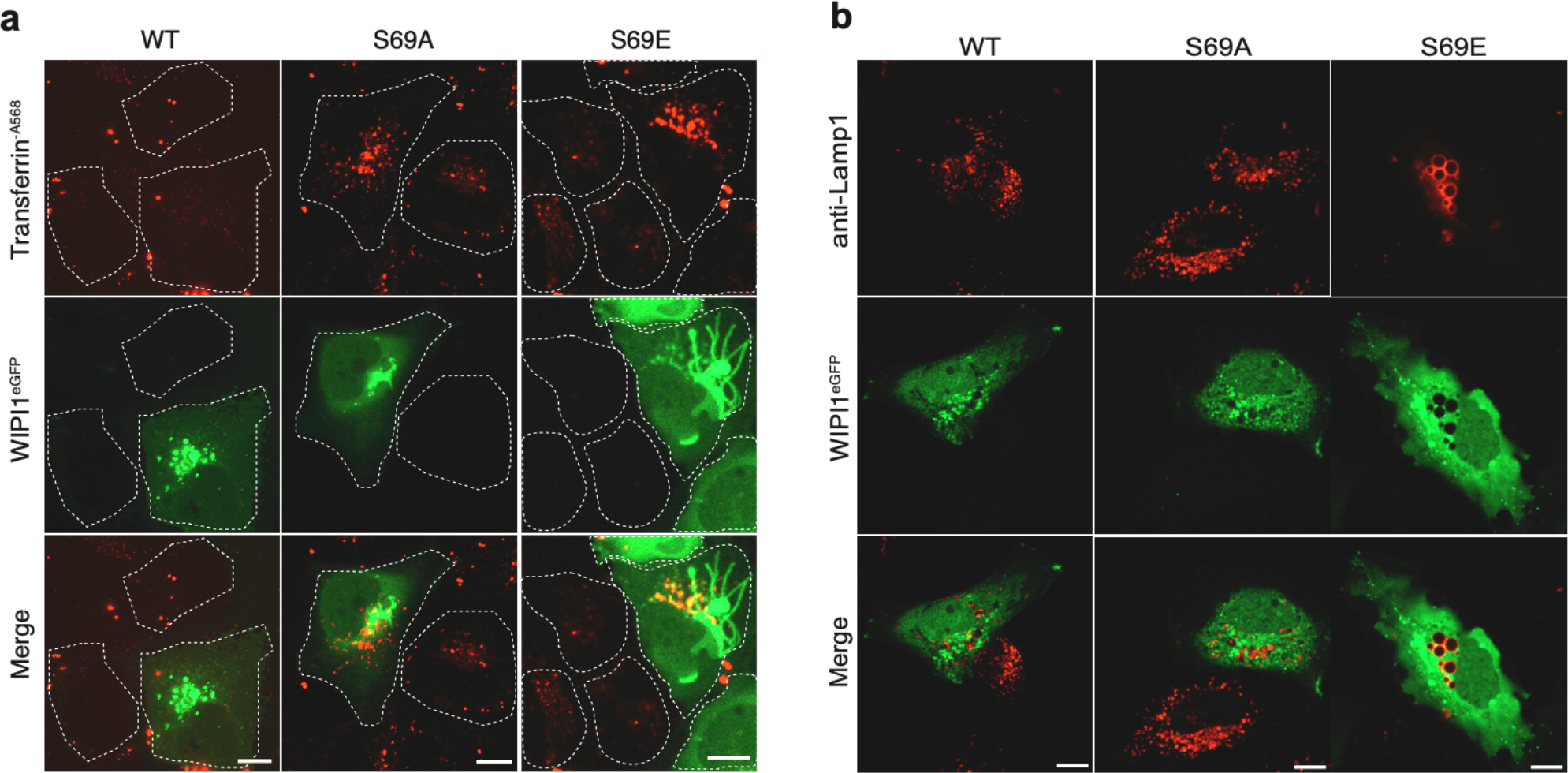
Dominant negative effect of EGFP-WIPI1^S69E^ on Transferrin recycling and on LAMP1 compartments. **a**, Tf recycling. HK2 cells were transfected with WIPI1^WT-EGFP^, WIPI1^S69E-EGFP^ or WIPI1^S69A-EGFP^ for 18 h. Then, they were serum-starved for 1h, loaded with Alexa Fluor 568-conjugated Tf, chased at 37°C for 1 h in medium without labeled Tf, and analyzed by confocal microscopy. Scale bar: 10 μM. The white dashed lines indicate the circumference of the cells. **b**, LAMP1 compartments. HK2 cells expressing WIPI1^WT-eGFP^, WIPI1^S69E-eGFP^ or WIPI1^S69A-eGFP^ were fixed 18 h after transfection. The cells were stained for immunofluorescence analysis with anti-LAMP1 antibody and imaged by confocal microscopy. Scale bars: 10 μm.

